# Alternative splicing downstream of EMT enhances phenotypic plasticity and malignant behavior in colon cancer

**DOI:** 10.1101/2022.03.01.482471

**Authors:** Tong Xu, Mathijs Verhagen, Rosalie Joosten, Wenjie Sun, Andrea Sacchetti, Leonel Munoz Sagredo, Véronique Orian-Rousseau, Riccardo Fodde

## Abstract

Phenotypic plasticity allows carcinoma cells to transiently acquire the quasi-mesenchymal features necessary to detach from the primary mass and proceed along the invasion-metastasis cascade. A broad spectrum of epigenetic mechanisms is likely to cause the epithelial–to-mesenchymal (EMT) and mesenchymal-to-epithelial (MET) transitions necessary to allow local dissemination and distant metastasis. Here, we report on the role played by alternative splicing (AS) in eliciting phenotypic plasticity in colon cancer.

By taking advantage of the identification of a subpopulation of quasi-mesenchymal and highly metastatic EpCAM^lo^ colon cancer cells, we here show that the differential expression of *ESRP1* and other RNA-binding proteins (RBPs) downstream of the EMT master regulator *ZEB1*, alters the AS pattern of a broad spectrum of targets including *CD44* and *NUMB*, thus resulting in the generation of specific isoforms functionally associated with increased invasion and metastasis. Additional functional and validation studies indicate that both the newly identified RBPs and the CD44s and NUMB2/4 splicing isoforms promote local invasion and distant metastasis and are associated with poor survival.

The systematic elucidation of the spectrum of EMT-related RBPs and AS targets in colon cancer and in other malignancies, apart from the insights in the mechanisms underlying phenotypic plasticity, will lead to the identification of novel and tumor-specific therapeutic targets.

## Introduction

Colon cancer still represents one of the major causes of cancer-related morbidity and mortality worldwide. Apart from its high incidence, the adenoma-carcinoma sequence along which colon cancer progresses has served as a classic model to elucidate the underlying genetic alterations representative of virtually all of the hallmarks of cancers^1^, possibly with the only exception of “activating invasion and metastasis”. As also reported in other epithelial cancers, the several steps of the invasion-metastasis cascade are not caused by genetic alterations but rather by transient morphological and gene expression changes of epigenetic nature^2, 3^. In this context, epithelial–mesenchymal transition (EMT), and its reverse MET, likely to represent the main mechanisms underlying local dissemination and distant metastasis^4, 5^. EMT is triggered at the invasive front of the primary colon carcinoma in cells earmarked by nuclear β-catenin and enhanced Wnt signaling, as the result of their physical and paracrine interactions with the microenvironment^6^. The acquisition of quasi-mesenchymal features allows local invasion and dissemination through the surrounding stromal compartment. Of note, EMT/MET should not be regarded as binary processes in view of previous reports highlighting the existence of metastable hybrid E/M states (partial EMT or pEMT) endowed with phenotypic plasticity and likely to underlie the reversible morphological and functional transitions necessary to successfully complete the invasion-metastasis cascade^7^.

The molecular basis of the epigenetic changes underlying EMT and MET is likely to encompass a broad spectrum of mechanisms ranging from chromatin remodeling and histone modifications to promoter DNA methylation, non-coding RNAs (e.g. micro RNAs), and alternative splicing (AS). The inclusion/exclusion of specific exons in mature mRNAs results in different protein isoforms with distinct biological functions. AS occurs in 92–94% of human genes leading to enriched protein density^8, 9^. Several sequence-specific RNA-binding proteins (RBPs) have been identified which bind pre-mRNAs to control AS in context-dependent fashion^10^. Multiple cancer-specific AS variants have been found to underlie progression and metastasis^11^. Likewise, alternative splicing has been suggested to play key roles in EMT/MET^12, 13^ and phenotypic plasticity^14^ in cancer by expression changes in RBP-encoding genes and their consequences for the modulation of downstream AS targets.

The *ESRP1* (epithelial splicing regulatory protein 1) gene encodes for an epithelial-specific RBP and splicing regulator shown to play a central role in EMT by modulating AS of EMT-associated genes including *FGFR2*, Mena, *CD44* and p120-catenin^4^. Relevant to the present study, ESRP1 was reported to regulate the EMT transition from CD44v (variable) to CD44s (standard) isoforms in breast and lung cancer progression^15, 16^. As for colon cancer, whether ESRP1 regulates alternative splicing of CD44 and other target genes downstream of EMT/MET activation during invasion and metastasis, is yet poorly understood.

Recently, we identified and thoroughly characterized subpopulations of CD44^hi^/EpCAM^lo^ colon cancer cells (here referred to as EpCAM^lo^) that coexist with their epithelial counterparts (CD44^hi^/EpCAM^hi^; for brevity EpCAM^hi^) through stochastic state transitions governed by phenotypic plasticity and pEMT^17^. Accordingly, EpCAM^lo^ cells feature highly invasive and metastatic capacities. Here, we took advantage of these *in vitro* models of phenotypic plasticity to study differential gene expression changes at upstream RBPs and AS variations at target genes. Among the top AS targets, CD44 and NUMB were selected for validation and functional studies in view of their known splicing isoforms and roles in stemness and cancer. Moreover, we provide an extensive list of additional EMT- and colon cancer-related RBPs and AS targets and show that the same CD44s and NUMB2/4 isoforms are conserved in ovarian and cervical cancer, i.e. independently of the distinct modalities through which these malignant cells metastasize.

## Materials and Methods

### Cell Cultures

The human colon cancer cell lines HCT116 and SW480, obtained from the European Collection of Authenticated Cell Culture (ECACC), were cultured in DMEM (11965092, Thermo Fisher Scientific) with 10% FBS (Thermo Fisher Scientific), 1% penicillin/streptomycin (Thermo Fisher Scientific, 15140122), and 1% glutamine (Gibco, 25030024), in humidified atmosphere at 37°C with 5% CO2.

### Plasmid transfection and lentiviral transduction

Stable transfection of the *ESRP1* (Sino Biological plasmid # HG13708-UT), *CD44s*, *CD44v6*, and NUMB1-4 (from V.O.R.) expression plasmids was performed using FuGENE HD transfection reagent (Promega, E2311) according to the manufacturer’s protocol and selected with Geneticin (Gibco, 10131035). As for the knockdown constructs, the *ESRP1*-shRNA plasmid (Horizon, V3THS_335722) was packaged by pPAX2 (Addgene # 12260) and pMD2.G (Addgene # 12259) into HEK293T. The virus-containing supernatant was collected 24 hrs. after transfection, filtered, and employed to infect the HCT116 and SW480 cell line. Selection was applied with 750 ng/ml puromycin (Invivogen, San Diego, USA) or 800 μg/ml of Geneticin selection for 1-2 weeks. The efficiency of overexpression and knockdown was assessed by qPCR and western blot 48-72 h after transfection.

### qRT-PCR and PCR analyses

Total RNA was isolated using TRIzol reagent (Thermo Fisher Scientific, 15596018) and was reverse-transcribed using high-capacity cDNA reverse transcription kit (Life Technologies, 4368814), according to the manufacturer’s instructions. qRT-PCR was performed using the Fast SYBR Green Master Mix (Thermo Fisher Scientific) on an Applied Biosystems StepOne Plus Real-Time Thermal Cycling Research with three replicates per group. Relative gene expression was determined by normalizing the expression of each target gene to GAPDH. Results were analyzed using the 2-(ΔΔCt) method. To validate isoform switches by RT-PCR, CD44-specific primers were as listed in Supplementary Table S3.

### Western analysis

Cells were lysed in 2X Laemmli buffer containing 4% SDS, 48% Tris 0.5M pH6.8, 20% glycerol, 18% H2O, bromophenol blue and 10% 1M DTT, and subjected to sodium dodecyl sulfate (SDS)-polyacrylamide gel electrophoresis (PAGE), followed by transfer onto polyvinylidene fluoride (PVDF) membranes (Bio-Rad). After blocking with 5% milk in TBS-Tween, the membranes were incubated with primary antibodies against ZEB1 (1:1000, Cell Signaling, #3396), ESRP1 (1:1000, Invitrogen, PA5-11520), CD44s (1:100, Invitrogen, MA5-13890), CD44v6 (1:1000, Abcam, VFF-7), NUMB (1:1000, Cell Signaling, C29G11) and β-actin (1:2000, Cell Signaling, 8547), followed by polyclonal goat anti-mouse/ rabbit immunoglobulins horseradish peroxidase (HRP)-conjugated secondary antibody (Dako) at appropriate dilutions. The signals were detected with Pierce ECT western blotting subtrade (Thermo) using Amersham AI600 (GE Healthcare, USA).

### Flow cytometry analysis and sorting

Single-cell suspensions generated in PBS supplemented with 1% FBS were incubated with anti-EpCAM-FITC (1:20, Genetex), and anti-CD44-APC (1:20, BD Pharmingen) antibodies for 30 min on ice and analyzed on a FACSAria III Cell Sorter (BD Biosciences). CD44^hi^EpCAM^hi^and CD44^hi^EpCAM^lo^ HCT116 and SW480 cells were sorted and cultured in humidified atmosphere at 37°C with 5% CO2 for 3-5 days before collecting RNA or protein, as previously described^17^. The subpopulation of cells mapping in between the CD44^hi^EpCAM^hi^ and CD44^hi^EpCAM^lo^ gates was labeled as intermediate and was further not employed for analysis.

### MTT assay

For MTT assay, 2×10^3^ HCT116, SW480 parental, CD44v6, CD44s, and NUMB1-4 OE cells were plated in 96 well plates and incubated at 37°C, 5% CO2. 24 hours later, in the culture medium was supplemented with 100μl 0.45 mg/mL MTT (3-(4,5-dimethylthiazol-2-yl)-2,5-diphenyltetrazolium bromide; Sigma-Aldrich) and again incubated for 3 hrs.. The 96-well plates were then centrifuged at 1,000 rpm for 5 min and the culture medium removed. MTT formazan precipitates were solubilized with DMSO. OD reading was performed at 595 nm with microplate reader (Model 550, Bio-Rad). Background measurements were subtracted from each data point. Experiments were performed in duplicate for each individual cell line and drug. Cell numbers were calculated every 24 hrs. for a 6 days period for proliferation analysis.

### Cell migration assay

Migration assay were conducted with 8-μm pore PET transwell inserts (BD Falcon™) and TC-treated multi-well cell culture plate (BD Falcon™). 5×10^4^ cells were seeded in 100 μl of serum-free growth medium in the top chamber. Growth medium containing 10% FBS was used as a chemoattractant in the lower chamber. After 24 hrs., cells migrated to the lower chamber were fixed with 4% PFA, stained with 0.1% trypan blue solution, and counted under the microscope.

### Mouse spleen transplantation

All mice experiment were implemented according to the Code of Practice for Animal Experiment in Cancer Research from the Netherlands Inspectorate for Health Protections, Commodities and Veterinary Public Health. Mice were fed in the Erasmus MC animal facility (EDC). NOD.Cg-Prkdc^scid^ Il2rg^tm1Wjl^/SzJ (NSG) mice from 8 to12 week-old were used for spleen transplantation. Anesthetics Ketamine (Ketalin®, 0.12 mg/ml) and xylazine (Rompun®, 0.61mg/ml) were given intraperitoneally, while the analgesic Carpofen (Rimadyl®, 5 mg/ml) was injected subcutaneously. 5x10^4^ HCT116 and SW480 cells resuspended in 50 μl PBS were injected into the exposed spleen with an insulin syringe and left for 15 minutes before splenectomy. Transplanted mice were sacrificed after 4 and 8 weeks and analyzed for the presence of liver metastases.

### Alternative splicing analysis

The following public available RNASeq (SRA database) data relative to RBP (RNA binding protein) knock-down (KD) studies were used: ESRP1-KD and RMB47-KD in the human non-small cell lung cancer cell line H358^18^ with accession ID SRP066789 and SRP066793; ESRP2-KD in the human prostate adenocarcinoma cancer cell line LNCaP^19^ with accession ID SRP191570; the QKI-KD in the oral squamous cell carcinoma cell line CAL27 datasets with accession number SRX8772405. Together with our own EpCAM^hi/lo^ RNASeq data obtained from the colon cancer cell lines^17^, the sequencing reads were mapped to GRCh37.p13.genome by STAR^20^ (https://www.gencodegenes.org/human/release_19.html). MISO^21^ was used to quantify AS events with annotation from https://miso.readthedocs.io/en/fastmiso/index.html#iso-centric. The MISO uses the alternative exon reads and adjacent conservative reads to measure the percentage of transcript isoform with specific exon included, termed Percentage Spliced In (PSI or Ψ). The PSI ranges from 0 (i.e. no isoform includes a specific alternative exon) to 1 (i.e. all of the isoforms detected comprise the alternative exon).

We removed alternative events with low expression of related transcript isoforms if less than 3 samples in a dataset had more than 10 informative reads to calculate the PSI. Next, we compared the PSI between RBPs KD and wild type in each cell line, as well as the PSI between EpCAM^hi^ and EpCAM^lo^ groups in the SW480 and HCT116 colon cancer cell lines. AS events were defined as differentially spliced events when the difference of mean PSI between two groups (Δpsi; differential Percentage Spliced In) was >10%.

### RNAseq analysis

RNA quality was first evaluated by NanoDrop and further purified by DNAse treatment followed by the TURBO DNA-free Kit protocol (Invitrogen). Samples were sequenced with the DNA nanoball (DNB) seq protocol (BGI) to a depth of 50 million reads per sample. Adapter sequences and low-quality sequences were filtered from the data using SOAPnuke software (BGI). Reads were aligned to the human reference genome build hg19 with the RNAseq aligner STAR (v2.7.9a) and the Homo sapiens GENCODE v35 annotation. Duplicates were marked with Sambamba (0.8.0) and raw counts were summed using FeatureCounts (subread 2.0.3). Downstream analysis was performed in R using the DESeq2 package (v1.30.1). After variance stabilizing transformation, principal component analysis was performed on each cell line separately. Differentially expressed genes were identified by comparing the different groups of ectopically expressing CD44 samples with a Wald test, and by selecting the genes with absolute log fold change above 1.5 and padj < 0.1. Gene set enrichment analysis was performed with the Fsgsea package using the HallMark geneset from the molecular signature database, and by selecting significant pathways based on normalized enrichment score (NES) > 1 and pvalue < 0.05.

### RNAseq data from primary (patient-derived) colon cancers

Patient data from The Cancer Genome Atlas (TCGA), with annotation of the consensus molecular subtypes (CMS) as described in Guinney et al.^22^ were integrated with splicing data from the TCGA splicing variant database (TSVdb, www.tsvdb.com). For splicing analysis, RNA-seq by expectation maximization (RSEM) values were log transformed and expression levels of each isoform (CD44std: isoform_uc001mvx, CD44v6: exon_chr11.35226059.35226187, NUMB1: isoform_uc001xny, NUMB2: isoform_uc001xoa, NUMB3: isoform_uc001xnz, NUMB4: isoform_uc001xob) were annotated to the patients. Isoform expression was compared in groups based on the CMS groups and tumor expression levels (*ZEB1*, *ESRP1*). Tumors were stratified on *ZEB1* expression levels using a log rank test top optimize overall survival differences (thresholds: 8.3, 8.6). Next, ESRP1 expression was used to purify the groups into ZEB1^hi^ESRP1^lo^ and ZEB1^lo^ESRP1^hi^ (thresholds: 11.6, 11.8). Survival analysis was done using the Kaplan-Meier method with the survival and survminer packages in R. Correlation analysis was done by computing the Pearson Correlation between the isoforms and whole gene expression levels as processed in Guinney et al.^22^. Likewise, association between isoform expression and pathway activity was evaluated by computing the Pearson Correlation between the isoforms and the average scaled expression values of the pathways, as defined in the HallMark gene set from the molecular signature database^23^.

### Data accessibility

The RNA-sequencing data from this study have been submitted to the Gene Expression Omnibus (GEO) database under the accession number GSE192877. Other data referenced in this study are publicly available and can be accessed from the GEO using GSE154927^17^, GSE154730 and Synapse using identifier syn2623706^22^ .

## Results

### Differential expression of RNA-binding proteins in the quasi-mesenchymal and highly metastatic EpCAM^lo^ colon cancer cells affects alternative splicing of a broad spectrum of downstream target genes

As previously reported, the EpCAM^lo^ subpopulation of colon cancer cells is earmarked by increased expression of the *ZEB1* transcription factor, responsible for EMT activation and for their quasi-mesenchymal and highly metastatic phenotype^17^. It has been established that in breast and pancreatic cancer *ZEB1*-driven EMT downregulates the expression of the RNA-binding protein and splicing regulator *ESRP1* as part of a self-enforcing feedback loop^24^. Accordingly, among the top differentially expressed genes between EpCAM^lo^ and EpCAM^hi^ in SW480 and HCT116 colon cancer cells, *ESRP1* was found to be downregulated both at the RNA and protein level in the quasi-mesenchymal subpopulation where *ZEB1* expression is upregulated (Figure 1A-C). Gain- and loss-of-function analyses of both genes confirmed the inter-dependence of their expression levels in both cell lines (Figure 1D-E). Of note, *ESRP1*-overexpression in the HCT116 and SW480 cell lines resulted in the dramatic reduction of their EpCAM^lo^ subpopulations and the expansion of the epithelial bulk, as shown by FACS analysis (Figure 1F, Suppl. Figure 1A). However, *ESRP1* knockdown (KD) only marginally affected CD44 though not EpCAM expression levels (Suppl. Figure 1B). The latter suggests that additional RNA binding proteins are likely to be involved in the alternative splicing regulation of the EpCAM^lo^ colon cancer subpopulation. Indeed, by taking advantage of the RBPDB database^25^, we found that, apart from *ESRP1*, consistent differential expression in the quasi-mesenchymal subpopulation of both cell lines was observed for *ESRP2*, *RBM47*, *MBNL3* (down-regulated) and *NOVA2*, *MBNL2*, (up-regulated). Other RBPs were found to be differentially expressed though in only one of the two cell lines (Suppl. Figure 1C). In validation of the clinical relevance of the RBPs found to be differential expressed between the EpCAM^hi/lo^ subpopulations derived from the SW480 and HCT116 cell lines, the RBP-coding genes *QKI*, *RBM24*, and *MBNL2* (up in EpCAM^lo^), and *ESRP1/2* and *RBM47* (down in EpCAM^lo^) were found to be respectively up- and down-regulated in the consensus molecular subtype 4 (CMS4) of colon cancers, responsible for ∼25% of the cases and earmarked by poor prognosis and a pronounced mesenchymal component (Suppl. Figure 1D)^22^.

**Figure 1.**
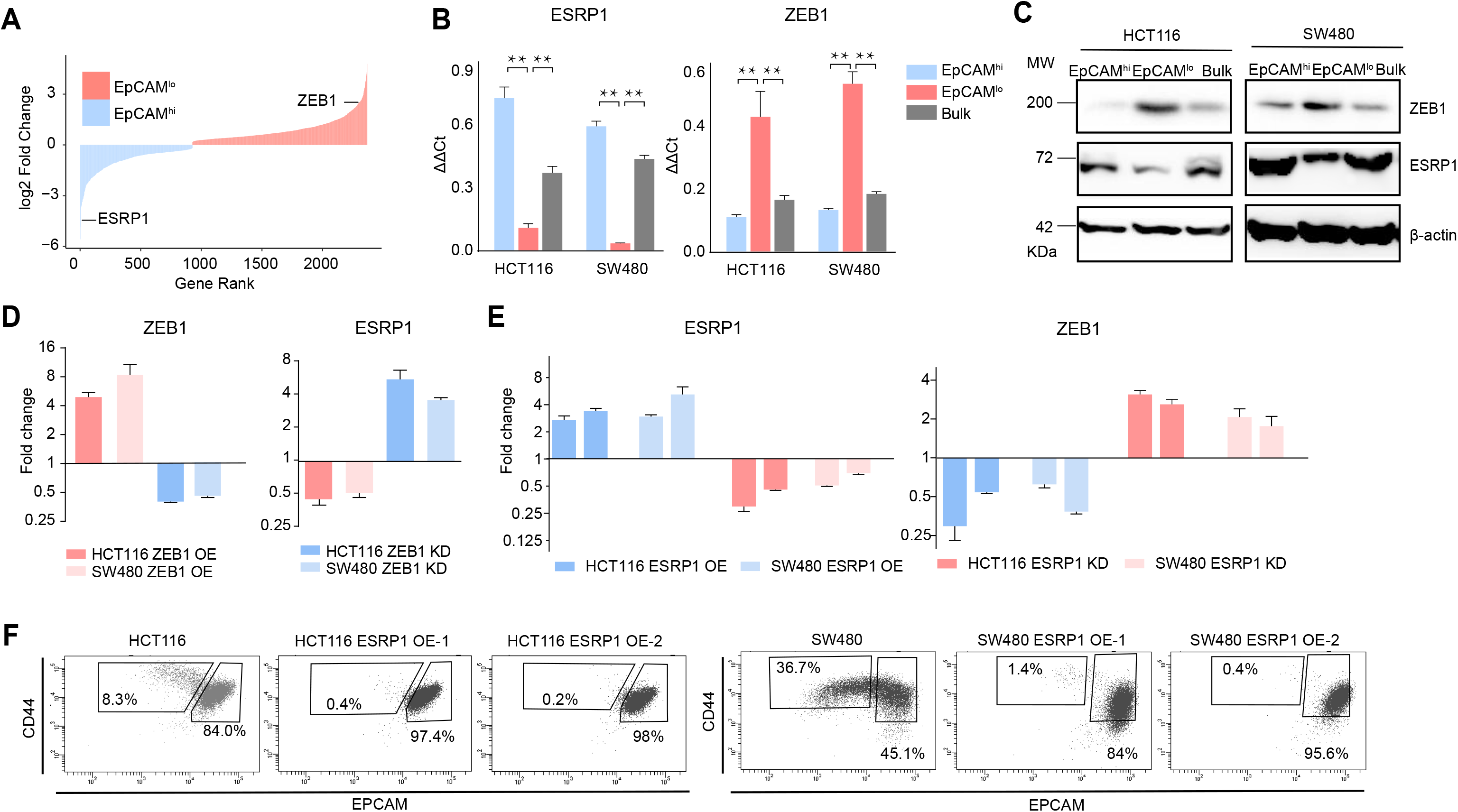
*ZEB1* and *ESRP1* differential expression in quasi-mesenchymal and highly metastatic EpCAM^lo^ colon cancer cells. **A.** Gene rank plot showing differentially expressed genes between EpCAM^hi^ and EpCAM^lo^ with combined analysis of HCT116 and SW480. **B.** RT-qPCR *ESRP1*- and *ZEB1* expression analysis of HCT116 and SW480 EpCAM^hi^, EpCAM^lo^, and bulk subpopulations. *GAPDH* expression was used as control (Means±SEM, n=3). ** = p<0.01. **C.** ESRP1 and ZEB1 western analysis in HCT116 and SW480 EpCAM^hi^, EpCAM^lo^, and bulk fractions. β-actin was used as loading control. **D.** RT-qPCR *ZEB1* and *ESRP1* expression analysis in *ZEB1*-OE and -KD HCT116 and SW480 cells. Expression values were normalized in each sample with those from the parental HCT116 and SW480 cell lines. HCT116 and SW480 cells transduced with the sh*ZEB1* lentivirus were induced by 1 µg/mL doxycycline for 72 hrs.. Expression values were normalized with those from non-induced cells; *GAPDH* expression was employed as control (Means±SEM, n=3). **E.** RT-qPCR *ZEB1* and *ESRP1* expression analysis in *ESRP1*-OE and -KD HCT116 and SW480 cells. Two independent *ESRP1*-OE clones were selected for each cell line. Expression values were normalized in each sample with those from the parental HCT116 and SW480 cell lines. HCT116 and SW480 cells transduced with the sh*ESRP1* lentivirus were induced by 1 µg/mL doxycycline for 72 hrs. Two independent clones were selected for each cell line. Expression values were normalized with those from non-induced cells; *GAPDH* expression was employed as control (Means±SEM, n=3). **F.** CD44/EpCAM FACS analysis of HCT116 and SW480 EpCAM^lo^ and EpCAM^hi^ subpopulations in ESRP1-OE cells. Two independent clones are showed for each cell lines.

Differentially spliced target genes between EpCAM^lo^ and EpCAM^hi^ colon cancer cells from the SW480 and HCT116 cell lines were selected based on exon skip splicing events with ΔPSI (differential Percentage Spliced In) values > 10%. The PSI value ranges from 0 to 1 and is a measurement of the percentage of isoform with an alternative exon included^26^. This resulted in a large and rather heterogeneous group of alternative spliced targets (n=1495; Suppl. Table 1a) with no clear enrichment in any specific gene ontology class (data not shown). In order to identify differentially spliced target genes in RBP-specific fashion, we took advantage of RNAseq data sets from previous *ESRP1*-, *ESRP2*-, *RBM47*-, and *QKI*-knockdown studies in different cancer cell lines and compared them with our own AS data relative to the EpCAM^hi/lo^ colon cancer subpopulations^17^ (Figure 2A and Suppl. Figure 2). A total of 32 common skipped exons events in 20 genes were identified between EpCAM^lo^ colon (both cell lines) and *ESRP1* KD H358 lung cancer cells^18^ (Figure 2A). More extensive lists of common *ESRP1* AS events and target genes were obtained when the SW480 and HCT116 cell lines were individually compared with the lung cancer study (Suppl. Table 1b-c). As for the alternative splicing targets of RBPs other than *ESRP1*, based on the available RNAseq data from knockdown studies of *ESRP2* (in the LNCaP cell line^19^), *RBM47* (H358^18^), and *QKI* (CAL27; GEO Accession: GSM4677985), several common and unique genes were found (Suppl. Figure 2 and Suppl. Table 2). Notably, four EMT-related genes (*CTNND1*^27^, *LSR*^28^, *SLK*^29^, and *TCF7L2*^30^) were common to all RBP KD studies analyzed (Suppl. Figure 2).

**Figure 2.**
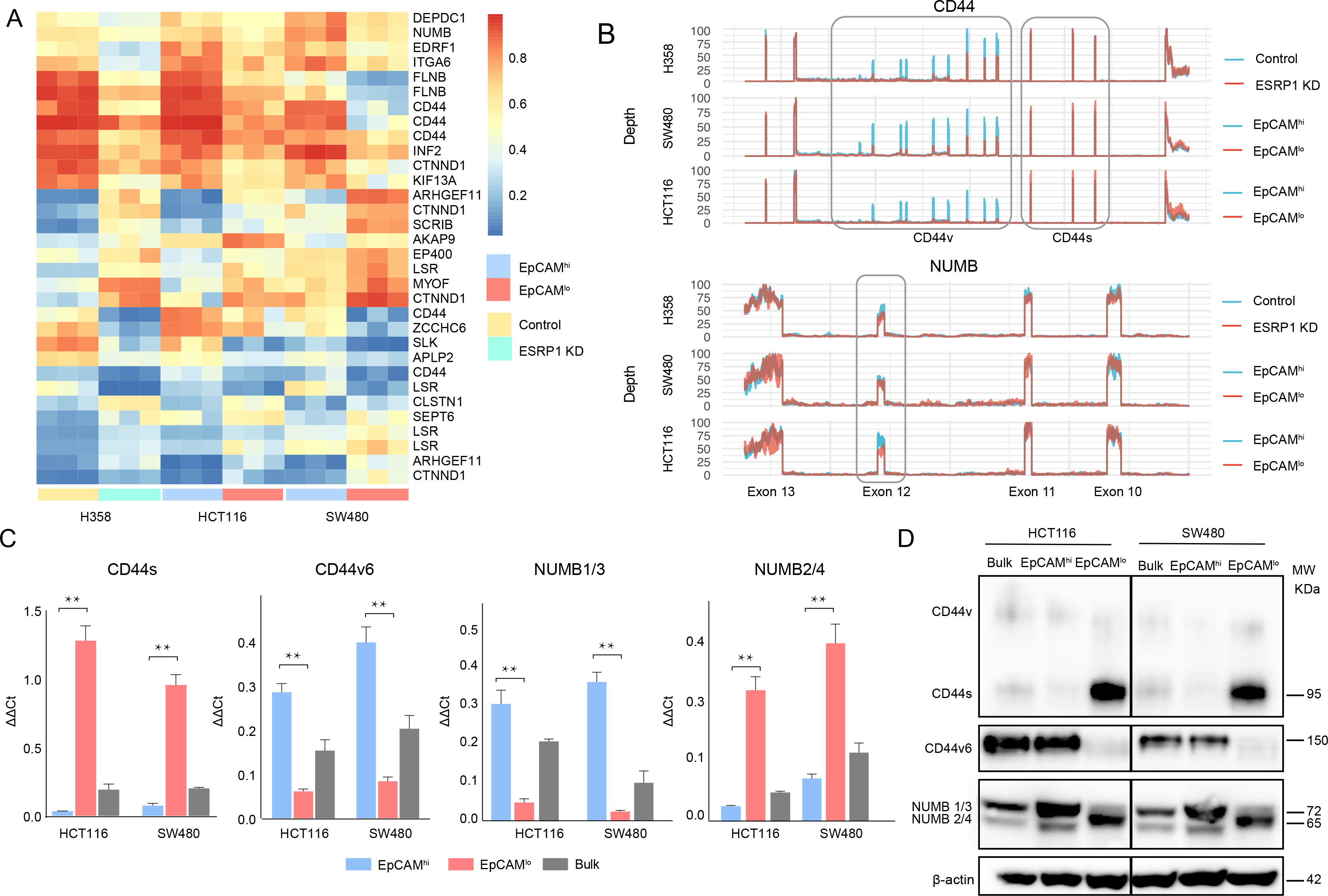
*ESRP1* downregulation in EpCAM^lo^ colon cancer cells affects alternative splicing of *CD44* and NUMB among a broad spectrum of downstream target genes. **A.** Heatmap of common AS events between RNAseq data from a previous *ESRP1*-KD study in human non-small cell lung cancer cells (H358)^18^ and our own HCT116 and SW480 EpCAM^hi^ and EpCAM^lo^ RNAseq data^17^. The gene list on the right of the heatmap encompasses AS variants earmarked by ΔPSI > 0.1. **B.** *CD44* and *NUMB* exon peak plots relative to the AS analysis of the RNAseq data obtained from a previous *ESRP1*-KD study in human non-small cell lung cancer cells (H358; upper graph)^18^ and from our own HCT116 (middle graph) and SW480 (lower graph) EpCAM^hi/lo^ analysis^17^. Each peak plot depicts the expression of specific exons; the height of each peak is indicative of the expression level of the specific exons. CD44v: CD44 exons v2 to v10. CD44v and CD44s, and NUMB exon 12 is highlighted by gray rectangles. **C.** RT-qPCR expression analysis of *CD44*s, *CD44*v6*, NUMB1*/*3* and *NUMB2*/*4* isoforms in HCT116 and SW480 EpCAM^hi^, EpCAM^lo^, and bulk subpopulations. *GAPDH* expression was employed as control (Means±SEM, n=3). ** = p<0.01. **D.** Western analysis of CD44s, CD44v6 and NUMB isoforms in HCT116 and SW480 EpCAM^hi^, EpCAM^lo^, and bulk subpopulations. β-actin was used as loading control.

### The CD44s and NUMB2/4 ESRP1-specific AS isoforms are preferentially expressed in EpCAM^lo^ colon cancer cells

From the newly generated lists of RBP-specific alternative splicing targets, we selected *CD44* and *NUMB* for further analysis, based both on their *ESRP1*-specific AS patterns and on their well-established roles in EMT, stemness/differentiation, and cancer progression.

CD44, a transmembrane cell surface glycoprotein, has been show to play key roles in inflammatory responses and in cancer metastasis^31^. The *CD44* gene encompasses 20 exons of which 1-5 and 16-20 are constant and exist in all isoforms. In contrast, exons 6-14, also referred to as variants exons v2-v10, are alternatively spliced and often deregulated in cancer^31^. The *NUMB* gene and its protein product have been involved in a broad spectrum of cellular phenotypes including cell fate decisions, maintenance of stem cell niches, asymmetric cell division, cell polarity, adhesion, and migration. In cancer, NUMB is a tumor suppressor that regulates, among others, Notch and Hedgehog signaling^32^. The mammalian *NUMB* gene encodes for 4 isoforms, ranging from 65 to 72 KD, differentially encompassing two key functional domains, i.e. the amino-terminal phosphotyrosine-binding (PTB) domain, and a C-terminal proline-rich region (PRR) domain^32^.

Based on the above ΔPSI-based AS analysis, decreased expression of CD44v (variable) isoforms was observed in EpCAM^lo^ and *ESRP1*-KD cells, accompanied by increased CD44s (standard) isoform expression (Figure 2B). Likewise, the NUMB2/4 isoforms appear to be preferentially expressed in EpCAM^lo^ and *ESRP1*-KD, accompanied by decreased NUMB1/3 expression (Figure 2B, Suppl Figure 2B). RTqPCR and western analyses validated these *in silico* data: CD44s and NUMB2/4 isoforms were preferentially expressed in EpCAM^lo^ colon cancer cells, in contrast with the increased CD44v and NUMB1/3 levels in EpCAM^hi^ cells (Figure 2C-D). In view of its previously suggested role in invasion and metastasis^33^, we focused on the CD44v6 isoform.

As reported above, AS events at the *NUMB* and *CD44* genes correlate with decreased ESRP1 expression. To confirm this observation, we up- and down-regulated *ESRP1* in the SW480 and HCT116 cell lines. The dox-inducible shRNA vector used for the KD studies reduces ESRP1 expression by 5-10 fold (Figure 1D-E) and resulted in the upregulation of the CD44s and NUMB2/4 isoforms at the mRNA and protein level in both cell lines (Figure 3A-B and Suppl. Figure 3). Likewise, *ESRP1* overexpression led to an increase in the CD44v6 and NUMB1/3 isoforms, found in association with the bulk of epithelial colon cancer cells (Figure 3C-D and Suppl. Figure 3).

**Figure 3.**
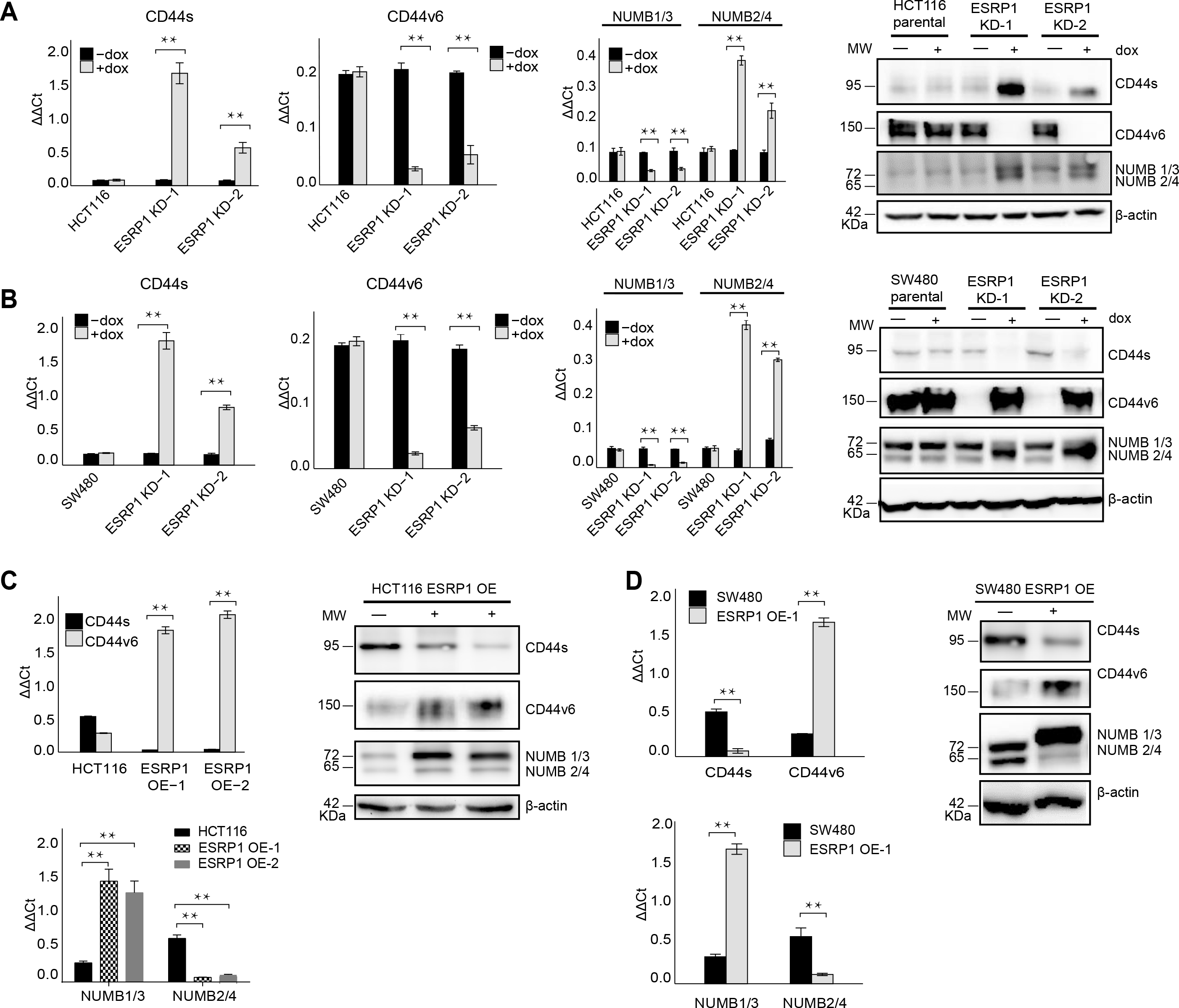
*ESRP1* differential expression regulates *CD44* and *NUMB* AS isoforms expression. **A.** RT-qPCR and western analysis of CD44s, CD44v6 and NUMB isoforms expression in *ESRP1*-KD (sh*ESRP1* transduced) HCT116 cells. Two independent HCT116 *ESRP1*-KD clones were employed. Cells were induced with 1 µg/mL doxycycline for 72 hrs. before analysis. *GAPDH* expression was employed as qRT-PCR control (Means±SEM, n=3). ** = p<0.01. β-actin was used as loading control for western blots. **B.** RT-qPCR and western analysis of CD44s, CD44v6 and NUMB isoforms expression in *ESRP1*-KD (sh*ESRP1* transduced) SW480 cells. Two independent SW480 *ESRP1*-KD clones were employed. Cells were induced with 1 µg/mL doxycycline for 72 hrs. before analysis. *GAPDH* expression was employed as qRT-PCR control (Means±SEM, n=3). ** = p<0.01. β-actin was used as loading control for western blots. **C.** RT-qPCR and western analysis of CD44s, CD44v6, and NUMB isoforms expression in *ESRP1*-OE HCT116 cells. Two independent HCT116 *ESRP1*-OE clones were employed. *GAPDH* expression was employed as qRT-PCR control (Means±SEM, n=3). ** = p<0.01. β-actin was used as loading control for western blots. **D.** RT-qPCR and western analysis of CD44s, CD44v6, and NUMB isoforms expression in *ESRP1*-OE SW480 cells. *GAPDH* expression was employed as qRT-PCR control (Means±SEM, n=3). ** = p<0.01. β-actin was used as loading control for western blots.

### Transcriptional and functional consequences of the CD44s and NUMB2/4 isoforms on colon cancer invasion and metastasis

In order to elucidate the functional contribution exerted by the newly identified CD44s and NUMB2/4 isoforms on the overall invasive and metastatic capacities of colon cancer cells, we first ectopically expressed each of them (individually and in combination for NUMB1/3 and 2/4) in the HCT116 and SW480 cell lines (Suppl. Figure 3E-H), and analyzed their consequences *in vitro* by cell proliferation, transwell migration assay, RTqPCR, western, FACS, and RNAseq, and *in vivo* by spleen transplantation. A significant increase in migratory capacity (Suppl. Figure 4A-B), comparable to that of EpCAM^lo^ cells sorted from the parental lines, was observed in SW480 and HCT116 upon overexpression of the CD44s and NUMB2/4 isoforms (Suppl. Figure 4A-B). Likewise, ectopic expression of the single NUMB2 or 4 isoforms resulted in increased migration rates when compared with NUMB1 and 3. In contrast, overexpression of CD44v6 and NUMB1/3, normally prevalent in the epithelial bulk (EpCAM^hi^) of both cell lines, did not affect their migratory properties (Suppl. Figure 4A-B).

In agreement with the migration assays, overexpression of CD44s and NUMB2/4 results in the significant upregulation of the EMT transcription factors (EMT-TFs) *ZEB1*, accompanied by the up- and downregulation regulation of mesenchymal and epithelial markers such as *VIM* (vimentin), *CDH1* (E-cadherin), and *EpCAM*, respectively (Suppl. Figure 4C). Of note, expression of *ESRP1*, the main upstream splicing regulator of both CD44 and NUMB, was also decreased in CD44s- and NUMB2/4-OE cells, in confirmation of the self-enforcing feedback loop that characterize its interaction with ZEB1 and EMT activation^24^. In agreement with the well-established regulation of Notch signaling by NUMB isoforms^32^, established Notch target genes and were accordingly up- (*HES1*, *HEY1*) and down-regulated (*ID2*) upon overexpression of NUMB2/4 (Suppl. Figure 4D).

FACS analysis was then employed to evaluate the overall effect of the ectopic expression of the specific CD44 and NUMB isoforms on the relative percentages of the EpCAM^hi/lo^ subpopulations in the HCT116 and SW480 cell lines. As shown in Figure 4A, CD44s overexpression led to a dramatic increase of the EpCAM^lo^ subpopulation at the expenses of EpCAM^hi^ cells. The opposite effect was observed with CD44v6, i.e. the enlargement of the EpCAM^hi^ gate and the corresponding decrease of EpCAM^lo^ cells. As for NUMB, ectopic expression of NUMB2/4 significantly increased the relative proportion of EpCAM^lo^ cells while reducing the size of the EpCAM^hi^ subpopulation, while the opposite was observed with NUMB1/3 (Figure 4B-C). Of note, the single NUMB2 and NUMB4 isoforms appear dominant in their capacity to enlarge the HCT116 and SW480 EpCAM^lo^ subpopulations, respectively. The same was true for NUMB1 and NUMB3 in the consequences of their ectopic expression in reducing the size of the HCT116 and SW480 EpCAM^lo^ fractions, respectively (Figure 4B-C). In agreement with the RTqPCR analysis of EMT markers, CD44s overexpression negatively affected overall proliferation rates in both cell lines, whereas the opposite was observed upon CD44v6 expression (Suppl. Figure 5A-B). Likewise, NUMB1/3 expression positively affected proliferation rates in HCT116 and SW480, whereas the NUMB2/4 isoforms exert the opposite effects. In both cases, synergistic effects were observed upon co-expression of NUMB1/3 and 2/4, when compared to the individual isoforms (Suppl. Figure 5C-D).

**Figure 4.**
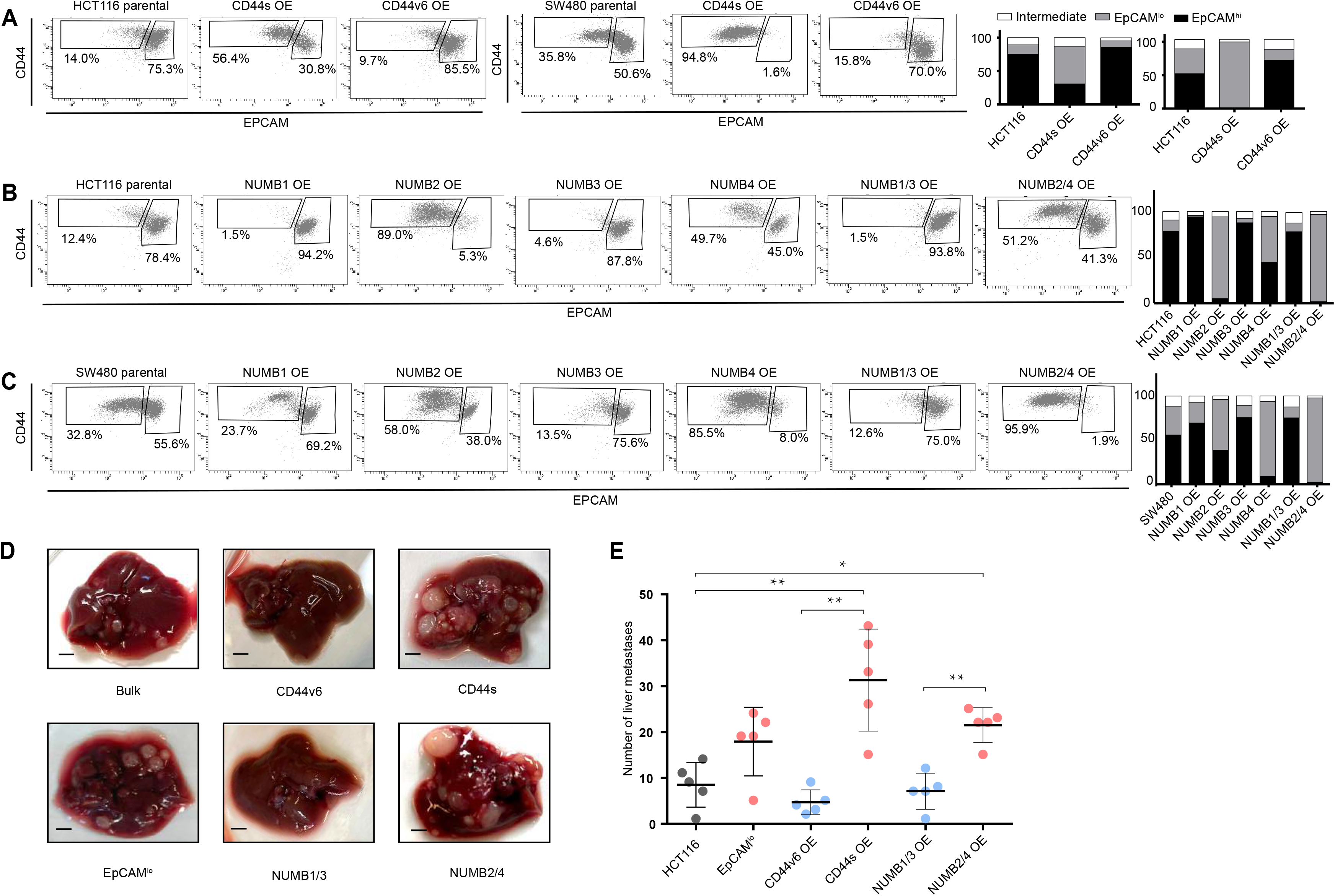
*CD44* and *NUMB* AS isoforms have opposite functions in quasi-mesenchymal and epithelial colon cancer cells and their capacity to metastasize the liver. **A.** CD44/EpCAM FACS analysis of EpCAM^lo^ and EpCAM^hi^ subpopulations in CD44s-OE (left) and CD44v6-OE HCT116 and SW480 cell lines. The bar charts on the right depict the percentages of EpCAM^lo^ and EpCAM^hi^ cells. The subpopulation of cells mapping in between, but yet outside, the CD44^hi^EpCAM^hi^ and CD44^hi^EpCAM^lo^ gates, is here labelled as ‘intermediate’. **B.** and **C.** CD44/EpCAM FACS analysis of EpCAM^lo^ and EpCAM^hi^ subpopulations in NUMB1 to 4-OE HCT116 and SW480 cells. The bar charts on the right depict the percentages of EpCAM^lo^ and EpCAM^hi^ cells. **D.** Macroscopic images of livers from mice spleen-injected with CD44s-, CD44v6-, NUMB2/4-, and NUMB1/3-OE HCT116 cells. HCT116 EpCAM^lo^ and bulk cells were used as positive control. Scale bar: 5 mm. **E.** Liver metastasis multiplicity after intrasplenic injection of CD44s-, CD44v6-, NUMB2/4-, and NUMB1/3-OE HCT116 cells. For each transplantation experiment, 5 × 10^4^ cells were injected in the spleen of recipient NSG mouse. Six weeks after injection, mice were sacrificed and individual tumors counted. * = p<0.05; ** = p<0.01.

In order to assess the *in vivo* the consequences of the ectopic expression of the CD44 and NUMB isoforms on the capacity of colon cancer cells to form metastatic lesions in the liver, parental HCT116 and SW480 cells and their CD44s-, CD44v6-, NUMB1/3-, and NUMB1/4-overexpressing counterparts were injected in the spleen of immune-incompetent recipient mice. In agreement with the *in vitro* results, overexpression of both NUMB2/4 and CD44s isoforms significantly increased the multiplicity of liver metastases, whereas CD44v6 and NUMB1/3 did not differ from the parental controls (Figure 4D-E).

Next, in order to elucidate the signaling pathways and molecular and cellular mechanisms triggered by the CD44 isoforms, we analyzed by RNAseq HCT116 and SW480 cells ectopically expressing CD44s and CD44v6. After dimension reduction with principal component analysis (PCA), the samples separated by group (i.e. CD44s-OE, CD44v6-OE, and controls) (Figure 5A). Notably, the CD44s-OE samples showed most distinct expression in both cell lines when compared to the parental and CD44v6-OE cell lines. In HCT116, the CD44v6 samples shared most similarity with the CD44s samples, while in SW480, the CD44v6 samples were most similar to the parental cell line. Thus, we observed both an isoform independent effect, presumably as the result of the ectopic CD44 expression (and most dominantly visible in HCT116), and an isoform dependent effect as depicted by the separation of CD44s and CD44v6 samples (Figure 5A). As expected, differential expression analysis of the CD44s and v6 isoforms overexpressing samples compared with the parental cell lines revealed an overall upregulation of gene expression (Suppl. Figure 6A). Next, in order to identify which genes are specifically upregulated by the different CD44 isoforms, we performed differential expression analysis between the CD44s samples and the CD44v6 samples. To this aim, we employed k-means clustering on the scaled expression values to separate genes specific for the CD44s isoform (e.g. *SPARC, ZEB1, VIM*), the CD44v6 isoform (e.g. *IL32, TACSTD2, CSF2*), and genes that were indiscriminative for the CD44v6 isoform or the parental cell lines (e.g. *MAL2, ESRP1, CDH1*) (Figure 5B). Finally, to identify the most distinct differences in signaling pathways and GO functional categories, we performed a gene set enrichment analysis (GSEA) by comparing the CD44s-with the CD44v6-overexpressing samples in the individual cell lines. Among the significantly altered pathways (normalized enrichment score > 1, pval < 0.05), epithelial mesenchymal transition (EMT) was the only one upregulated in CD44s vs. CD44v6 in both cell lines (Figure 5C-D). Additional pathways and GO categories activated by CD44s appeared to be cell line specific, e.g. Wnt beta catenin signaling (HCT116) and oxidative phosphorylation (SW480). Of note, the detailed GSEA analysis evidenced how several inflammatory (TNF/NFκB; IL6/JAK/STAT3; IFα/γ; ILK2/STAT5) and signaling (KRAS, MYC, E2F) pathways were common to both CD44s and v6, presumably as the result of the ectopic CD44 expression, regardless of the isoform (Suppl. Figure 6B).

**Figure 5.**
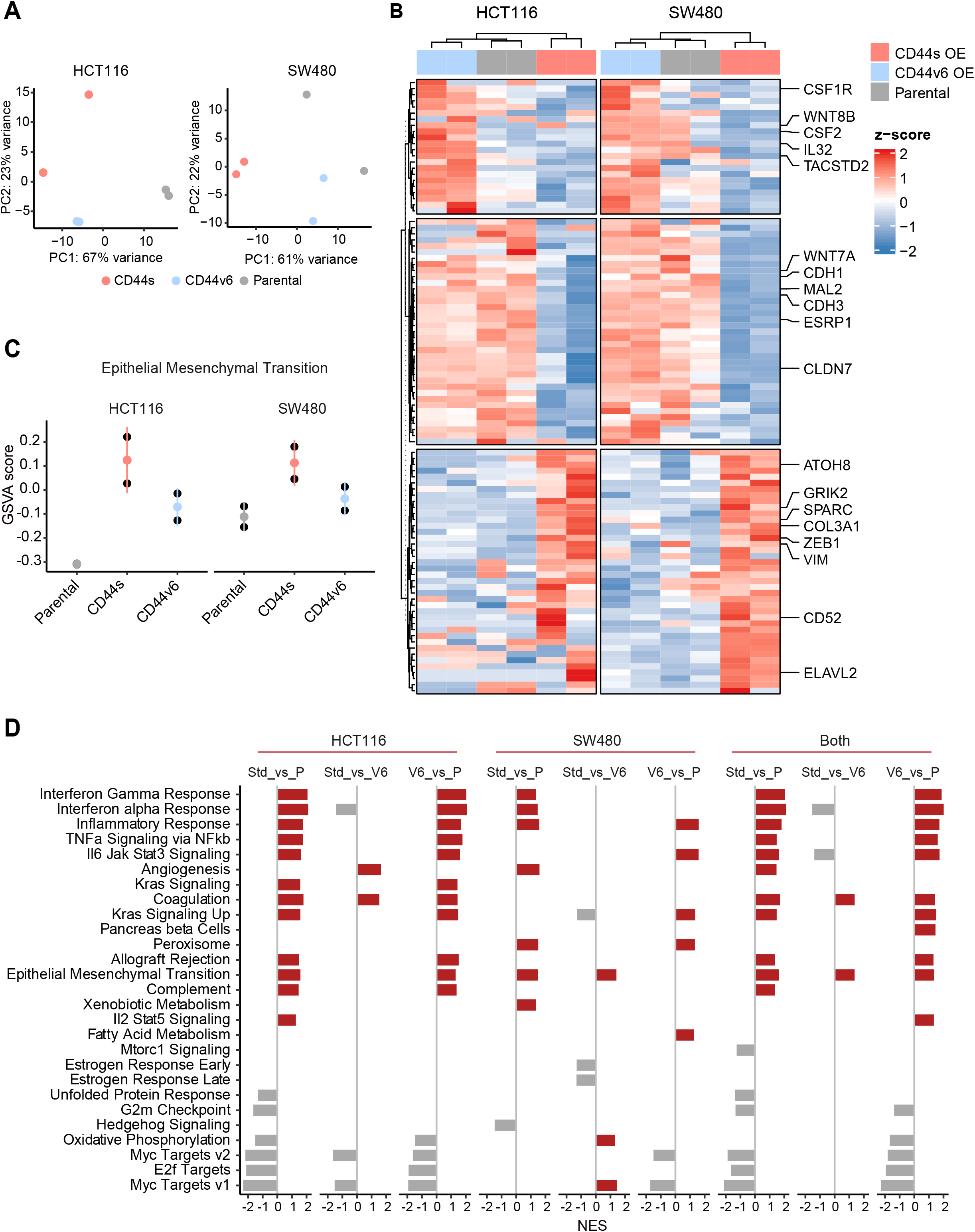
RNAseq analysis of CD44s- and CD44v6-expressing colon cancer cells reveals a broad spectrum of downstream AS targets and biological functions. **A.** Principal Component Analysis (PCA) of RNAseq profiles from CD44s- and CD44v6-OE HCT116 and SW480 cell lines. **B.** Heatmap of differentially expressed gene among HCT116 and SW480 CD44s-OE, CD44v6-OE, and parental cells. **C.** Gene Set Enrichment Analysis (GSEA) of epithelial mesenchymal transition (EMT) in expression profiles from HCT116 and SW480 parental, CD44s-OE, and CD44v6-OE cells. Normalized enrichment score (NES) > 1, and pval < 0.05. **D.** Gene Set Enrichment Analysis (GSEA) of HCT116 and SW480 expression profiles in parental, CD44s-OE, CD44v6-OE cells compared with each other. Plots show only significantly altered pathways, with normalized enrichment score (NES) > 1, and pval < 0.05.

### Increased ZEB1 and decreased ESRP1 expression correlate with the NUMB2/4 and CD44s isoforms and with poor overall survival

In order to assess the clinical relevance of the results obtained with the SW480 and HCT116 cell lines, we analyzed RNAseq data from patient-derived colon cancers available from the public domain and the scientific literature. To this aim, the TCGA Splicing Variants Database (TSVdb; www.tsvdb.com) was employed to integrate clinical follow-up data with RBP and AS expression profiles obtained from The Cancer Genome Atlas project (TCGA) and from the Guinney et al study^22^ on the classification of human colon cancers into four consensus molecular subtypes (CMS1–4). The main limitation of this approach is the low representation of quasi-mesenchymal (EpCAM^lo^-like) subpopulations in bulk RNAseq preparations and the masking effect that the majority of epithelial (EpCAM^hi^-like) cancer cells are likely to cause. To identify tumors enriched in EpCAM^lo^-like cells, we first stratified them based on *ZEB1* expression (*ZEB1*>8.6: ZEB1^hi^; ZEB1<8.3: ZEB1^lo^; 8.2<ZEB1<8.6: Intermediate). Subsequently, we used *ESPR1* expression levels to further define the tumors into *ZEB1*^hi^*ESRP1*^lo^ (*ESRP1*<11.8; hereafter referred to as *ZEB1*^hi^), *ZEB1*^lo^*ESRP1*^hi^ (*ESRP1*>11.6; hereafter referred to as *ZEB1*^lo^). Tumors with intermediate *ZEB1* expression levels and tumors with *ESRP1* expression levels outside these thresholds were defined as intermediate (Figure 6A). Kaplan-Meier analysis showed that *ZEB1*^hi^ tumors have an overall decreased survival probability (p = 0.045) (Figure 6B). Next, we compared the expression of CD44 and NUMB isoforms across the *ZEB1*^hi/lo^ tumors. Notably, while no significant differences were observed based on the expression level of the whole CD44 and NUMB genes, significant differences were found for their specific isoforms (Figure 6C). Analysis of the specific isoforms expression across the different consensus molecular subtypes^22^ revealed elevated CD44s and NUMB2/4 expression in the CMS4 subtype, known to be enriched in mesenchymal lineages in tumor and TME cells, and strongly associated with poor survival and the greatest propensity to form distant metastases (Figure 6D). Likewise, the majority of the *ZEB1*^hi^ group was composed of the CMS4 subtype (72%), while the *ZEB1*^lo^ group was mainly contributed by CMS2 (49%) and CMS3 tumors (31%), with few CMS4 tumors (1%) (Figure 6E).

**Figure 6.**
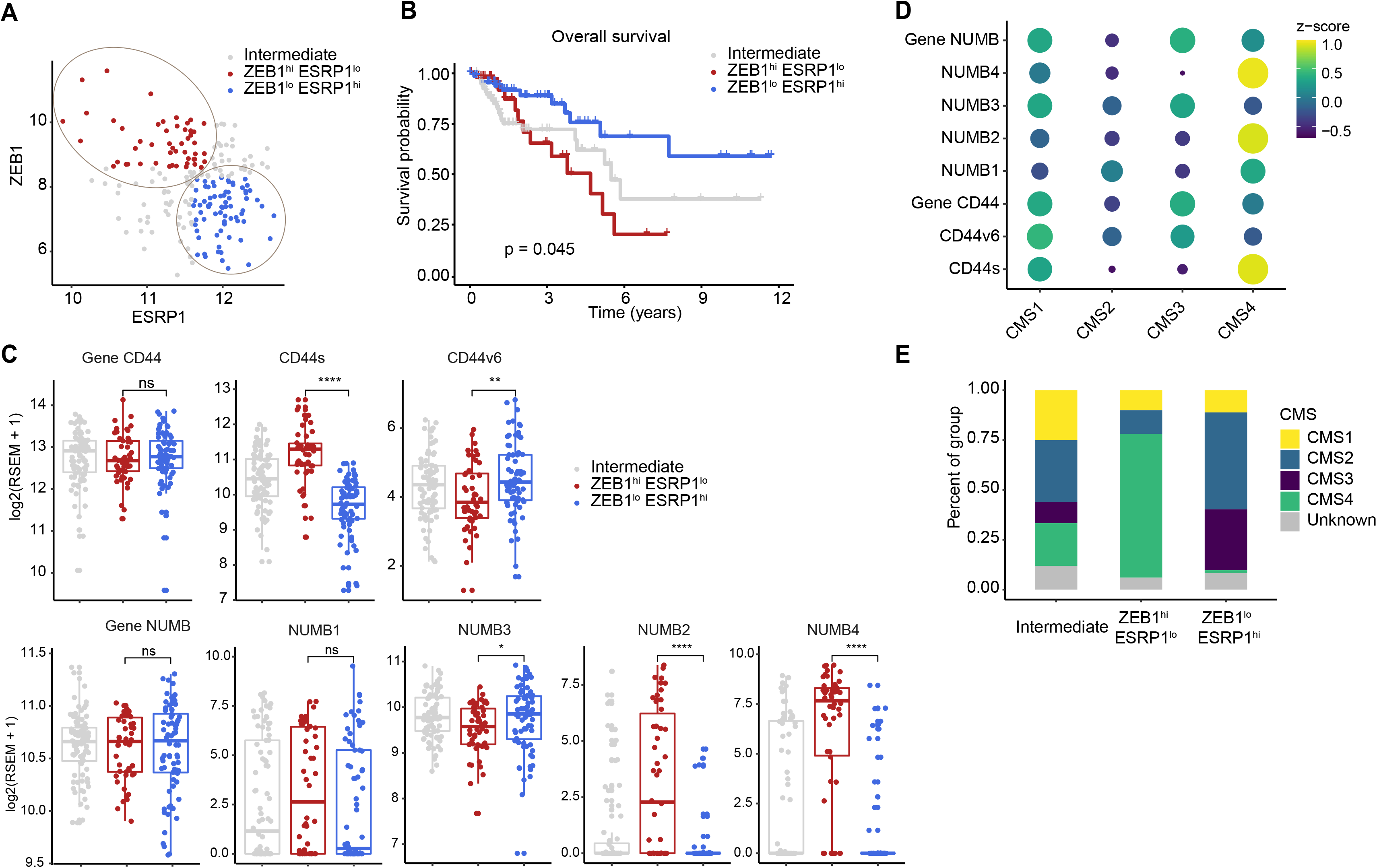
Increased *ZEB1* and decreased *ESRP1* expression correlate with the NUMB2/4 and CD44s isoforms and with poor overall survival. **A.** RNAseq data from the Cancer Genome Atlas (TCGA) were subdivided in 3 groups based on *ZEB1* and *ESRP1* expression level: *ZEB1*^hi^*ESRP1*^lo^ (*ZEB1*^hi^, red dots), *ZEB1*^lo^*ESRP1*^hi^ (*ZEB1*^lo^, blue dots), and intermediate (grey dots). **B.** Kaplan Meier analysis of overall survival in the *ZEB1*^hi^*ESRP1*^hi^ and *ZEB1*^lo^*ESRP1*^lo^ patient groups. **C.** Box plots showing CD44 and NUMB gene and isoforms expression across the *ZEB1*^hi^*ESRP1*^lo^, *ZEB1*^lo^*ESRP1*^hi^, and intermediate patient groups. **D.** Dot plot analysis of the z-score scaled expression values of CD44s, CD44v6, NUMB1-4 isoforms across the 4 colon cancer consensus molecular subtypes (CMS). **E.** Stacked bar plot showing the composition of the CMS subtypes across the *ZEB1*^hi/lo^ and intermediate patient groups.

Next, we correlated the expression of CD44s/v6 isoforms in patient-derived colon tumors with the differentially expressed genes (DEGs) identified in the isoform-overexpressing cell lines (Figure 7A). While overall *CD44* expression correlated with both isoforms, the DEGs from the CD44s-OE samples showed specific correlation with CD44s expression in patient-derived tumors (e.g. *SPARC, ZEB1*), the DEGs from the CD44v6 samples correlated with CD44v6 but not with CD44s (e.g. *KDF1, ESRP1*).

**Figure 7.**
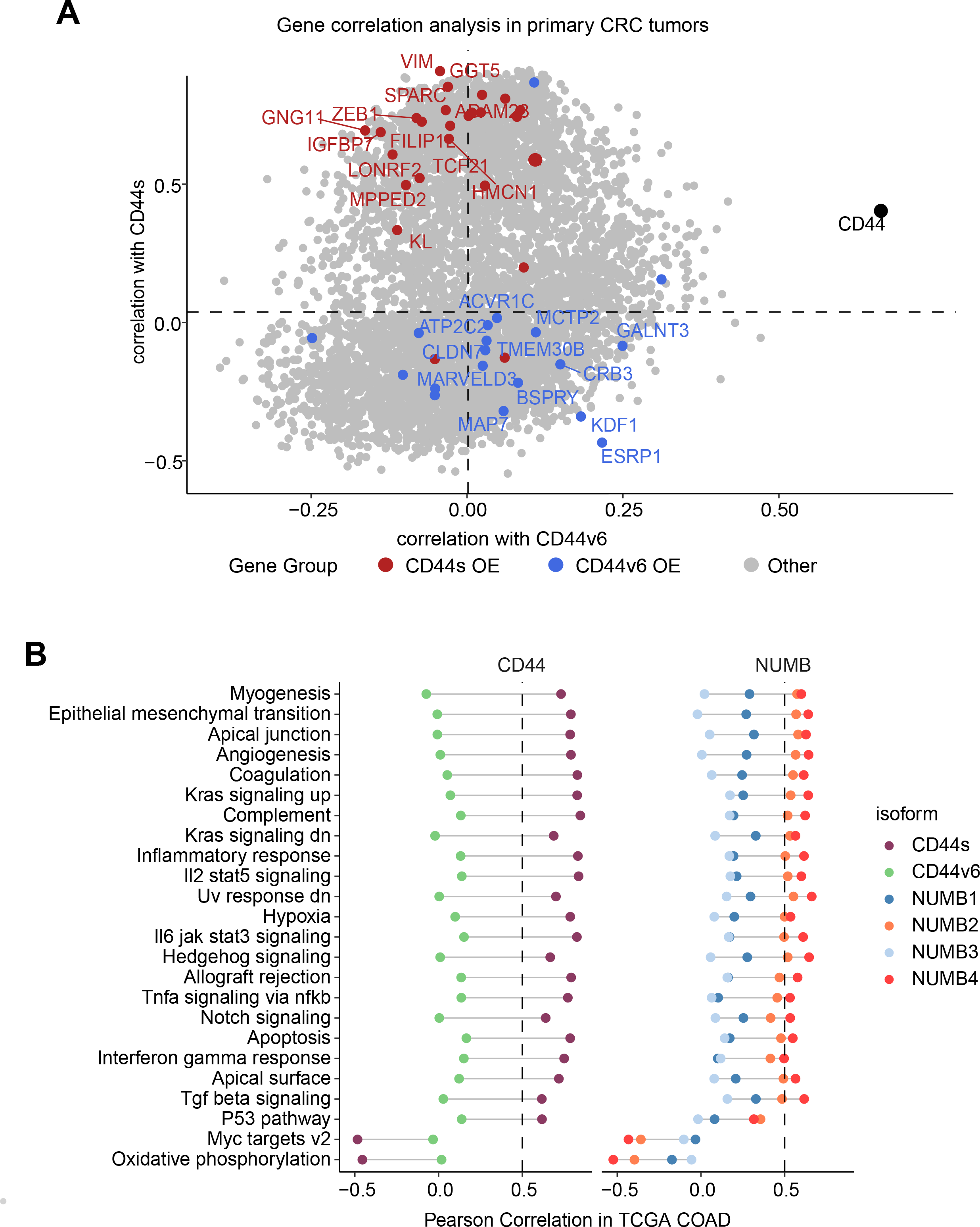
Gene and pathway correlation analyses of CD44 and NUMB isoforms in patient-derived colon cancers. **A.** Gene correlation analysis showing the correlation of gene expression with CD44s and CD44v6 isoform expression in the TCGA patient cohort. Differentially expressed genes from CD44s- (red) and CD44v6-OE (blue) RNAseq data are highlighted. **B.** Pathway correlation analysis showing the correlation of pathway activity CD44 and NUMB isoform expression in the TCGA patient cohort.

Last, we correlated the CD44 and NUMB isoforms expression in patient-derived colon cancers with functional signatures obtained by averaging the scaled expression levels for each of the hallmark sets^23^. The CD44s and NUMB2/4 isoforms showed overall similar correlating hallmarks and pathways. However, the same was not true when compared to the CD44v6- and NUMB1/3-associated functional signatures. Here, most invasion/metastasis-relevant hallmarks (e.g. EMT, angiogenesis, apical junctions) showed a positive correlation with CD44s and NUMB2/4, though not with CD44v6 and NUMB1/3 (Figure 7B).

In sum, we confirmed a switch in isoform expression (CD44v6 vs. CD44s and NUMB1/3 vs. NUMB2/4) as a function of *ESRP1* and *ZEB1* expression in colon cancer. Expression of the EpCAM^lo^–specific isoforms (CD44s and NUMB2/4) is elevated in CMS4 tumors overall survival.

### Upregulation of the NUMB2/4 and CD44s isoforms is common to quasi-mesenchymal cells from cancers other than colon

In order to assess whether the preferential expression of the NUMB2/4 and CD44s isoforms is specific to the modalities of local invasion and distant metastasis characteristic of colon cancer, we interrogated expression profiling data previously obtained by comparing epithelial and quasi-mesenchymal subpopulations from ovarian (OV90) and cervical (SKOV6) cancer cell lines (*manuscript in preparation*). Ovarian cancer, because of the distinct anatomical localization of the primary lesion, metastasizes the abdominal cavity with very different modalities than colon cancer, namely by peritoneal dissemination rather than local dissemination into the stroma microenvironment followed by intra- and extravasation of the portal blood stream^34, 35^. On the other hand, metastasis in carcinoma of the cervix occurs both by lymphatic or hematogenous spread to the lung, liver, and bones. We asked whether, notwithstanding the distinctive patterns of metastatic spread, the CD44s and NUMB2/4 isoforms were preferentially expressed in the corresponding EpCAM^lo^ RNAseq profiles. To this aim, EpCAM^hi/lo^ subpopulations from OV90 and SKOV6 were sorted and analyzed by RNAseq and RTqPCR, similar to our previous study on colon cancer^17^. As shown in Suppl. Figure 7, both NUMB2/4 and CD44s isoforms appear to be upregulated in the OV90 and SKOV6 cell lines, as also validated by RTqPCR.

## Discussion

The capacity to invade the tumor microenvironment and to form distant metastases undoubtedly represents the most clinically relevant hallmark of epithelial cancer cells. However, the complexity and diversity of the obstacles that carcinoma cells encounter along the invasion-metastasis cascade require transient and reversible changes that cannot be explained by the *de novo* acquisition of genetic alterations. Instead, epigenetic modifications underlie phenotypic plasticity, i.e. the capacity of cancer cells with a given genotype to acquire more than one phenotype in a context-dependent fashion^36^. Epithelial-to-mesenchymal and mesenchymal-to-epithelial transitions (EMT/MET) are central to the phenotypic plasticity characteristic of metastasizing carcinoma cells and are prompted by a broad spectrum of epigenetic mechanisms ranging from chromatin remodeling by histone modifications, DNA promoter methylation, non-coding RNAs, and alternative splicing (AS)^37^. Here, we have taken advantage of our previous identification of phenotypic plastic and highly metastatic EpCAM^lo^ colon cancer cells^17^ to characterize the genome-wide AS events that accompany EMT/MET state transitions between the epithelial bulk (EpCAM^hi^) and the quasi-mesenchymal subpopulation.

In view of the central role played by RNA-binding proteins in AS, we first identified RBP-coding genes differentially expressed between the EpCAM^lo^ and EpCAM^hi^ fractions of two commonly employed colon cancer cell lines, representative of the chromosomal- and microsatellite-instable subtypes (SW480, CIN; HCT116, MIN)^38^. The Epithelial Splicing Regulatory Protein 1 and 2 genes (*ESRP1*/*2*)^39^, the “*splicing masterminds*” of EMT^40, 41^, were found among the top downregulated RBP coding genes in EpCAM^lo^ colon cancer cells, as part of a self-enforcing feedback loop with the EMT-TF *ZEB1*^24^. Accordingly, *ZEB1* upregulation in EpCAM^lo^ colon cancer cells is invariably accompanied by *ESRP1/2* downregulation, and *ZEB1*^hi^/*ESRP1*^lo^ colon cancers, predominantly belonging to the mesenchymal CMS4 subgroup, have a significantly worse survival outcome when compared with *ZEB1*^lo^/*ESRP1*^hi^ patients.

Apart from *ESRP1*, several other RBP-coding genes were found to be differentially expressed between epithelial and quasi-mesenchymal colon cancer cells. Whereas the majority of RBP-coding DEGs, like *ESRP1*, appear to be downregulated upon EMT induction (*ESRP1*/*2*, *RBM14*/*19*/*47*, *MBNL3*, *HNRPAB*/*PF*, *USAF2*), others were activated in the quasi-mesenchymal EpCAM^lo^ fraction (*NOVA2*, *MBNL2*, *QKI*, *SRSF5*, *HNRNPH*, *RBM24*/*43*). Accordingly, in patient-derived colon cancers stratified according to their consensus molecular signature, the same *QKI*, *RBM24*, and *MBNL2* genes were found to have increased expression in CMS4 tumors, known for their pronounced mesenchymal composition and poor prognosis^22^. Of note, the mesenchymal nature of CMS4 tumors has previously been questioned as these lesions often feature pronounced infiltration from the surrounding microenvironment, the extent of which might cover their true cellular identity other than representing a mere contamination from the tumor microenvironment^42, 43^. As shown in our previous study^17^, the EpCAM^lo^ cells do represent *bona fide* quasi-mesenchymal colon cancer cells, enriched among CMS4 cases, and likely responsible for their poor prognosis. The observed upregulation of RBPs such as quaking (*QKI*) is caused by the presence in its 3’UTR of target sequences of the miR-200 family of microRNAs^44, 45^. The latter is analogous to the regulation of the expression of the EMT-TF *ZEB1* gene, whose activation during EMT is regulated by the same microRNA family^46^. Accordingly, the significantly reduced levels of all five miR-200 members in EpCAM^lo^ cells^17^ underlies the coordinated upregulation of both *ZEB1* and *QKI*.

The increased expression of other RBP-coding genes such as *RBM24* and *MBNL2* (muscleblind-like 2) in CMS4 tumors and in EpCAM^lo^ cells is also in sharp contradiction with their alleged tumor suppressing roles in colon and other cancers^47, 48^. Of note, MBNL2 regulates cancer migration and invasion through PI3K/AKT-mediated EMT^48^ and its overexpression in breast and cancer cell lines inhibits their metastatic potential^49^. In contrast to *MBNL2*, *MBNL3*, a distinct member of the muscleblind family, is downregulated in EpCAM^lo^ colon cancer cells, similar to what reported in prostate cancer by Lu and colleagues^50^. *NOVA2*, a member of the Nova family of neuron-specific RNA-binding proteins, was also upregulated in the quasi-mesenchymal cells from both cell lines, possibly as the result of the differential expression of miR-7-5p^51^, as previously shown in non-small cell lung^51^ and prostate^50^ cancer. The identification the AS targets downstream of specific RBPs in quasi-mesenchymal cancer cells from different malignancies will likely clarify these apparent contradictions and shed light the functional roles of distinct members of the splicing machinery in EMT and metastasis.

The spectrum of AS target genes downstream of the RBPs differentially expressed in EpCAM^lo^ colon cancer cells appears extremely broad when it comes to specific cellular processes or signaling pathways. Nonetheless, comparison of our RNAseq data with KD studies of specific RBPs from the public domain (*ESRP1*/*2*^19^, *RBM47*^18^, and *QKI* [GEO Accession: GSM4677985]) allowed us to identify common and unique AS target genes associated with specific downstream effectors. By following this admittedly imperfect approach, the top 4 AS targets common to all of the above-mentioned RBPs notwithstanding their up- or downregulation in EpCAM^lo^ colon cancer cells, i.e. *CTNND1* (δ- or p120-catenin), *LSR* (Lipolysis Stimulated Lipoprotein Receptor), *SLK* (STE20 Like Kinase), and *TCF7L2* (Transcription Factor 7-Like 2, or TCF4) are known regulators and effectors of epithelial-to-mesenchymal transition^27–30^, thus pointing to the central role played by alternative splicing in the regulation of EMT in the malignant evolution of colon cancer.

Here, we have focused on CD44 and NUMB as two ESRP1-specific AS target genes with well-established functional roles in EMT and in cancer invasion and metastasis. The CD44s and NUMB2/4 isoforms appear to be specifically expressed in quasi-mesenchymal colon cancer cells both from the immortalized cell lines and from patient-derived tumors, with a striking enrichment in the CMS4 subgroup of colon cancer patients. In contrast, the CD44v6 and NUMB1/3 isoforms are preferentially expressed in the epithelial bulk of the tumor. The latter, as far as CD44v6 is concerned, sharply contrasts what previously reported by Todaro et al.^33^ where this specific isoform was found to earmark the colon cancer stem cells (CSCs) which underlie metastasis. CD44v6 and other ‘variable’ CD44 isoforms (CD44v4-10) earmark *Lgr5*^+^ intestinal stem cells (ISCs), i.e. the stem cells of origin of intestinal tumors, and accordingly promote adenoma formation *in vivo*^52–54^. A plausible explanation for the discordant results lies in the epithelial nature of the models employed in the above study and in the requirement of both EMT and MET for the completion of the invasion-metastasis cascade^5^. By employing tumor spheres and freshly sorted CD133^+^ tumor cells, Todaro et al. focused on epithelial CSCs where, as observed in normal ISCs, the CD44v6 isoform is predominantly expressed, and is necessary for EMT to occur upon interaction with c-MET^33^. The CD44v6 isoform is strictly required for c-MET activation by hepatocyte growth factor (HGF, or scatter factor)^55^ and as such plays an essential role in triggering EMT at the invasive front where tumor cells are exposed to these TME-secreted factors. Our own immunoprecipitation studies confirmed that CD44v6 but not CD44s binds to cMET in response to HGF stimulation (*data not shown*). Therefore, HGF/SF stimulation of colon cancer cells along the invasive front will trigger the acquisition of quasi-mesenchymal characteristics and the AS-driven switch from CD44v6 to CD44s, the latter unable to bind HGF and as such controlling the extension of EMT activation. The reverse switch will take place upon the activation of the mesenchymal-to-epithelial transitions necessary for the colonization of the distal metastatic site. From this perspective, both CD44 isoforms are essential for the completion of the invasion-metastasis cascade. The functional relevance of the CD44s isoforms has been highlighted in malignancies other than colon cancer, namely in prostate^50^ and breast cancer where it activates, among others, PDGFRβ/Stat3 and Akt signaling to promote EMT and CSC traits^15, 56^. GO analysis of the RNAseq profiles from colon cancer cells ectopically expressing CD44s highlighted a broader spectrum of signaling pathways likely to underlie EMT. Accordingly, analysis of RNAseq data from primary colon cancers stratified for their CD44s expression revealed an equally broad spectrum of downstream EMT-related biological processes. Of note, among the DEGs identified upon CD44s ectopic expression which correlate with *ZEB1*^hi^/*ESRP1*^lo^ (and CMS4) colon cancers, the *SPARC* gene, a partial EMT marker in the EpCAM^hi/lo^ state transitions^17^, was found.

Expression of NUMB2/4 isoforms both in cells lines and in patient-derived colon tumors is associated with signaling pathways and GO categories largely overlapping with those linked to CD44s (and CD44v6 with NUMB1/3), possibly suggesting synergism between AS at these genes. Accordingly, NUMB is involved in a broad spectrum of cellular phenotypes in homeostasis and in cancer where it mainly function as a tumor suppressor^32^. NUMB inhibits EMT by suppressing the Notch signaling pathway. As such, downregulation of NUMB can induce an EMT phenotype in isoform-specific fashion. Analysis of colon cancer cells individually overexpressing each of the four isoforms revealed an increased basal Notch signaling in NUMB2 and 4, as shown by the expression of the ‘universal’ targets *HES1* and *HEY1*. Instead, ectopic expression of NUMB1/3 resulted in increased transcriptional levels of the more atypical Notch signaling target *ID2*. Although the functional consequences of the NUMB2/4 (and 1/3) isoforms on Notch regulation of EMT is yet unclear, it seems plausible that the complex network of AS targets activated downstream the RBP-coding DEGs, including CD44, NUMB and many others as shown here, will eventually lead to the ‘just-right’ level of plasticity needed to allow both the ‘mesenchymalization’ during local invasion and systemic dissemination, and the reacquisition of epithelial features at the distant site of metastasis. Overall, it appears that alternative splicing substantially contribute to the epigenetic mechanisms that underlie EMT/MET in colon cancer metastasis. The systematic elucidation of the RBPs and AS targets will not only elucidate the cellular and molecular mechanisms underlying phenotypic plasticity as the most clinically relevant hallmark of cancer, but it will also offer novel tumor-specific targets for therapeutic intervention based on small molecule inhibitors and even RNA vaccination.

## Authors’ contribution

T.X. performed most of the experiments and wrote the first manuscript draft. M.V. contributed the *in silico* analysis and validation of the AS results in patient-derived RNAseq data. R.J. and A.S. contributed to the implementation of PCR, mouse, and FACS experiments. W.S. analysed AS splicing in the RNAseq data. L.M.S. and V.O-R. contributed to the CD44 AS analysis and critically revised the manuscript. RF conceived the experimental strategy and wrote the manuscript.

The authors declare no competing interests.

## Acknowledgements

We are grateful to Dr. Juan Valcarcel, (CRG, Barcelona, Spain), for his critical reading of the manuscript.

## Supplementary Figure Legends

**Figure S1.**
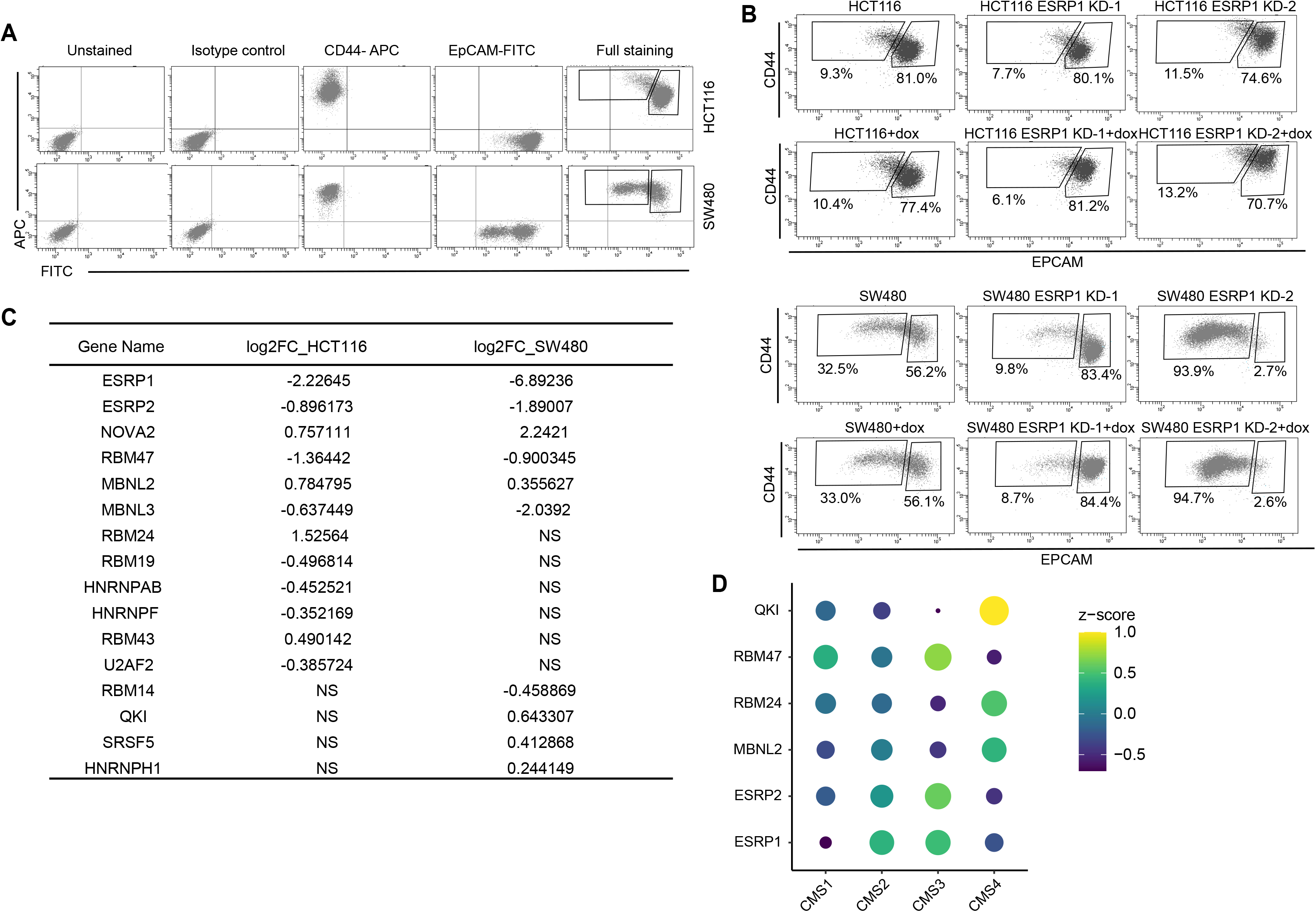
*ESRP1* and RBPs functional and expression analysis in cell lines and patient-derived colon cancers. **A.** FACS isotype and compensation controls in the analysis of the HCT116 and SW480 cell lines. The gates relative to the EpCAM^hi/lo^ subpopulations are specifically designed for the HCT116 and SW480 cell lines, as shown for the full staining. For the sake of simplicity and readability, the quadrants showing negative, single positive, and double positive regions, have not been repeated in the figures encompassing FACS analyses. **B.** CD44/EpCAM FACS analysis of EpCAM^lo^ and EpCAM^hi^ subpopulations in ESRP1-KD (*shESRP1*-transduced) HCT116 and SW480 cells. Cells were induced with 1 µg/mL doxycycline for 72 hrs. before analysis. **C.** List of RBPs differentially expressed between EpCAM^lo^ and EpCAM^hi^ subpopulation in SW480 and HCT116. The RBPs list was from reference (10). **D.** Dot plot analysis of the z-score scaled RBPs’ expression values across the 4 colon cancer consensus molecular subtypes (CMS; annotated according to Guinney et al.^20^. RNAseq data were obtained from the COAD (COlon-ADenoma) tumors of The Cancer Genome Atlas (TCGA) deposited in the TCGA Splicing Variants Database (TSVdb), (n=206 primary tumors).

**Figure S2.**
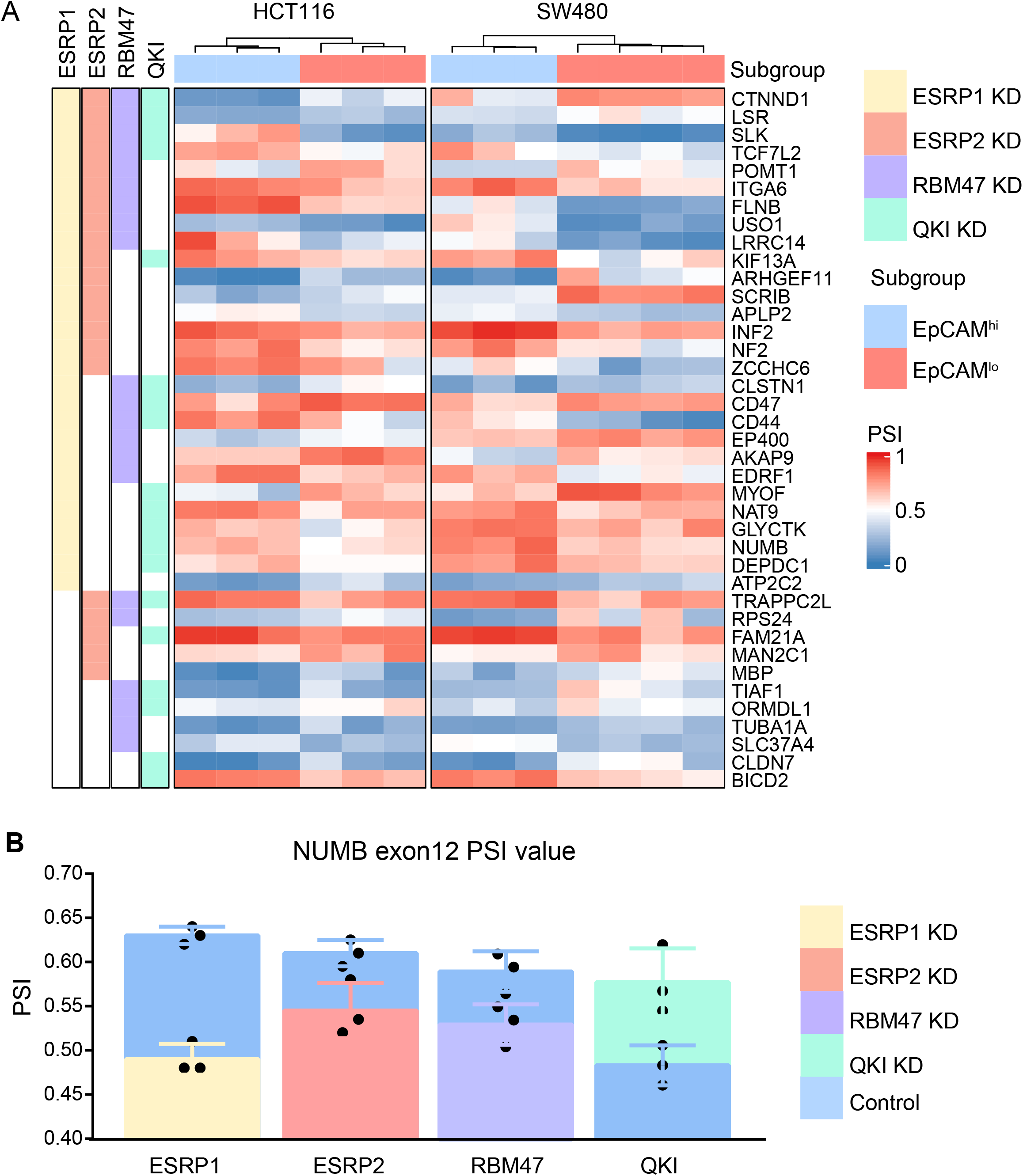
*ESRP1/2-, RBM47*-, and *QKI*-regulated AS targets. A. Heatmap of the alternative splicing events observed by comparing previously published RNAseq data from RBP-KD studies (ESRP1-KD in H358^18^, ESRP2-KD in LNCaP^19^, RBM47-KD in H358^18^, and QKI-KD in CAL27 [GEO Accession: GSM4677985]) with our own HCT116/SW480 EpCAM^hi/lo^ RNAseq data^17^. Shared AS targets between RBPs KD cells, and HCT116/SW480 EpCAM^hi/lo^ subpopulations are shown. The gene list on the right side of the heatmap encompasses variants earmarked by ΔPSI > 0.1. The colored bars on the left of the heatmap shows if there are variants spliced by different RPBs. Color in white means AS is not involved in. B. PSI value of NUMB exon12 between ESRP1 KD, ESRP2 KD, RBM47 KD, QKI KD and control cells.

**Figure S3.**
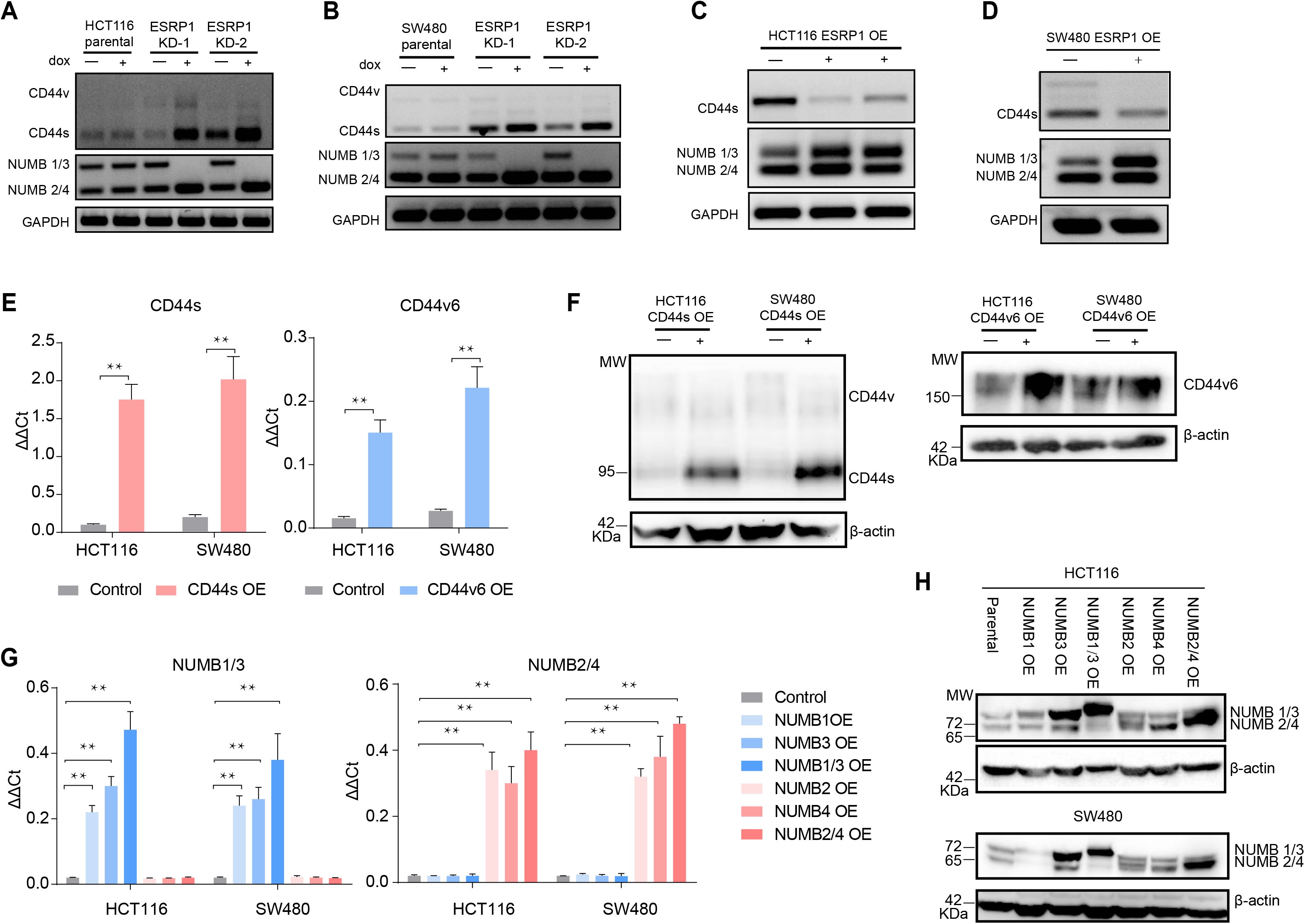
*ESRP1, CD44*, and *NUMB* isoforms analysis in over-expressing and knock-down colon cancer cell lines. RT-PCR analysis of CD44 and NUMB isoforms expression in HCT116 (**A**) and SW480 (**B**) ESRP1-KD (*shESRP1*-transduced) cells, and in HCT116 (**C**) and SW480 (**D**) *ESRP1*-OE cells. Cells were induced with 1 µg/mL doxycycline for 72 hrs. before RNA isolation. *GAPDH* was used as control. **E.** RT-PCR analysis of CD44s and CD44v6 expression in HCT116 and SW480 CD44s-(left), and CD44v6-OE (right) cells. *GAPDH* was used as control. (Means±SEM, n=3). ** = p<0.01. **F.** Western analysis of CD44s and CD44v6 expression in HCT116 and SW480 CD44s-(left), CD44v6-OE (right) cells. β-actin was used as loading control for western blots. **G.** RT-PCR analysis of NUMB1-4 isoforms expression in HCT116 and SW480 NUMB1-4 OE cells. *GAPDH* was used as control. (Means±SEM, n=3). ** = p<0.01. **H.** Western analysis of NUMB1-4 isoforms expression in HCT116 and SW480 NUMB1-4 OE cells. β-actin was used as loading control for western blots.

**Figure S4.**
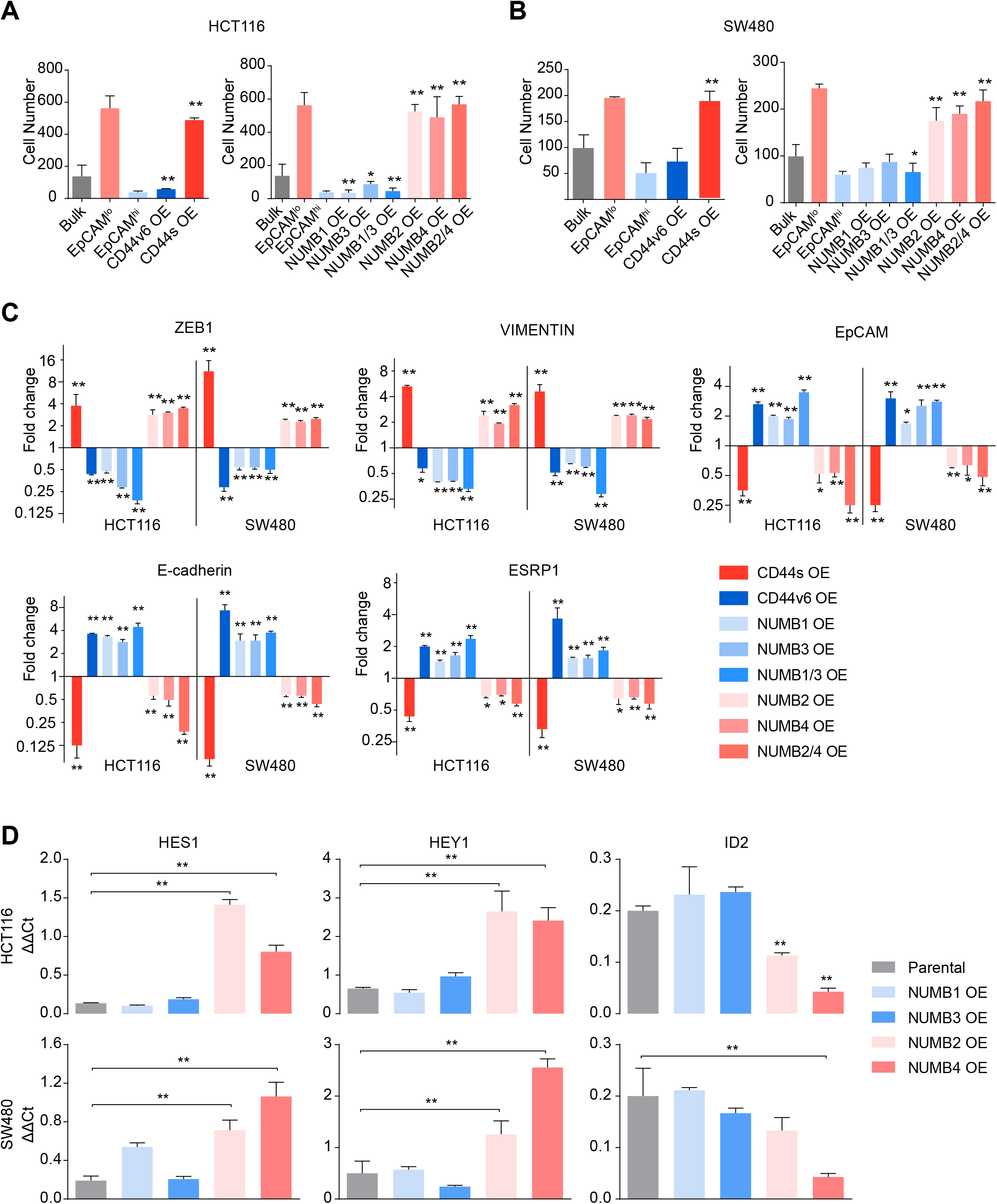
*CD44* and *NUMB* isoform-specific expression affects cell migration and Notch signaling activation. **A.** Migration assay analysis of HCT116 CD44s-, CD44v6-, and NUMB1/4-OE cells. EpCAM^lo^ and EpCAM^hi^ cells were used as controls. Each bar represents the mean ± SD of cells migrated to the bottom of the transwell from two independent experiments. **B.** Migration assay analysis of SW480 CD44s-, CD44v6-, and NUMB1/4-OE cells. EpCAM^lo^ and EpCAM^hi^ cells were used as controls. Each bar represents the mean ± SD of cells migrated to the bottom of the transwell from two independent experiments. **C.** RT-qPCR analysis of EMT-TFs in HCT116 and SW480 CD44s-, CD44v6-, and NUMB1/4-OE cells. *GAPDH* expression was used as control, normalized with the HCT116 or SW480 parental in each sample (Means±SEM, n=3). Increased gene expression is depicted by red bars, whereas downregulation – when compared with parental cells-is shown by blue bar. **D.** RT-qPCR analysis of the Notch signaling pathway markers *HES1*, *HEY1*, and *ID2* in HCT116 and SW480 NUMB1/4-OE cells. *GAPDH* expression was used as control (Means±SEM, n=3). * = p<0.05; ** = p<0.01.

**Figure S5.**
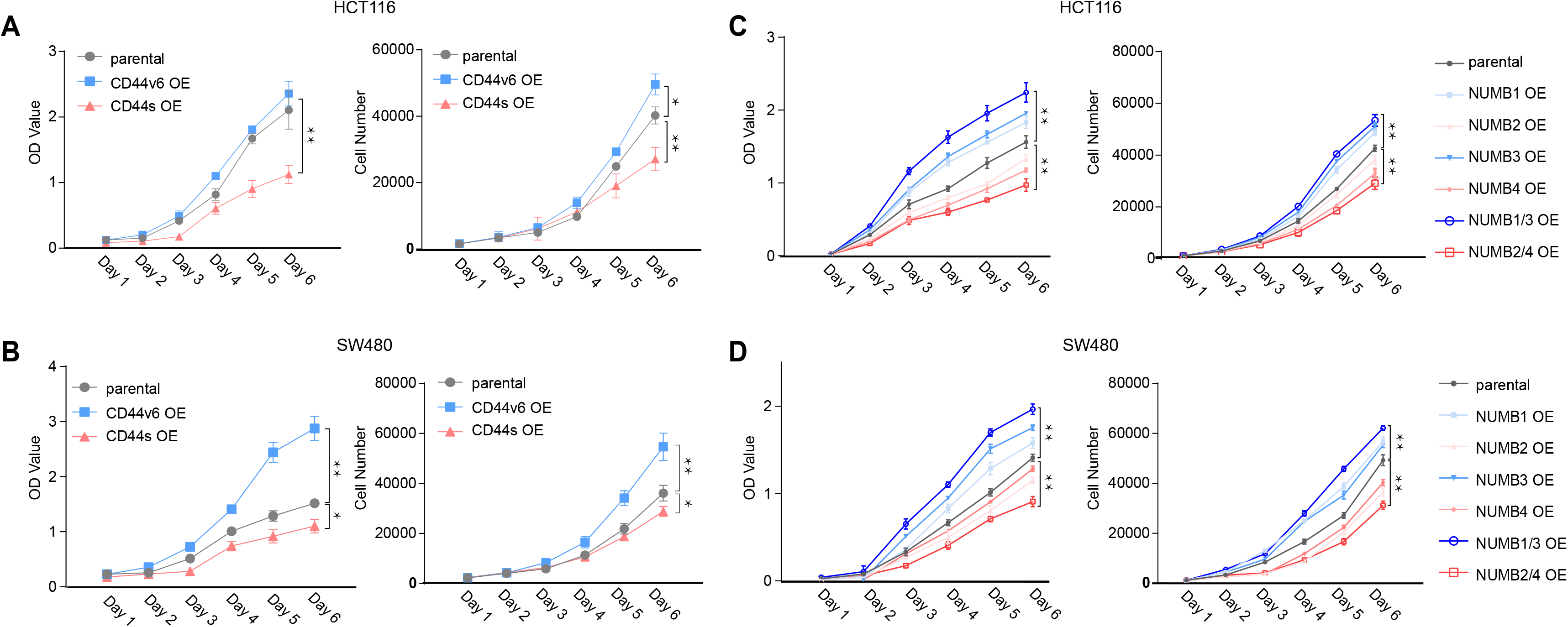
CD44 and NUMB isoforms regulate colon cancer cell proliferation. **A.** Proliferation assays of HCT116 CD44s- and CD44v6-OE cells. Both OD values and cell multiplicities are shown from day 1 to 6 (Means±SEM, n=3). * = p<0.05; ** = p<0.01. **B.** Proliferation assays of SW480 CD44s- and CD44v6-OE cells. Both OD values and cell multiplicities are shown from day 1 to 6 (Means±SEM, n=3). * = p<0.05; ** = p<0.01. **C.** Proliferation assays of HCT116 NUMB1/4-OE cells. Both OD values and cell multiplicities are shown from day 1 to 6 (Means±SEM, n=3). ** = p<0.01. **D.** Proliferation assays of SW480 NUMB1/4-OE cells. Both OD values and cell multiplicities are shown from day 1 to 6 (Means±SEM, n=3). ** = p<0.01.

**Figure S6.**
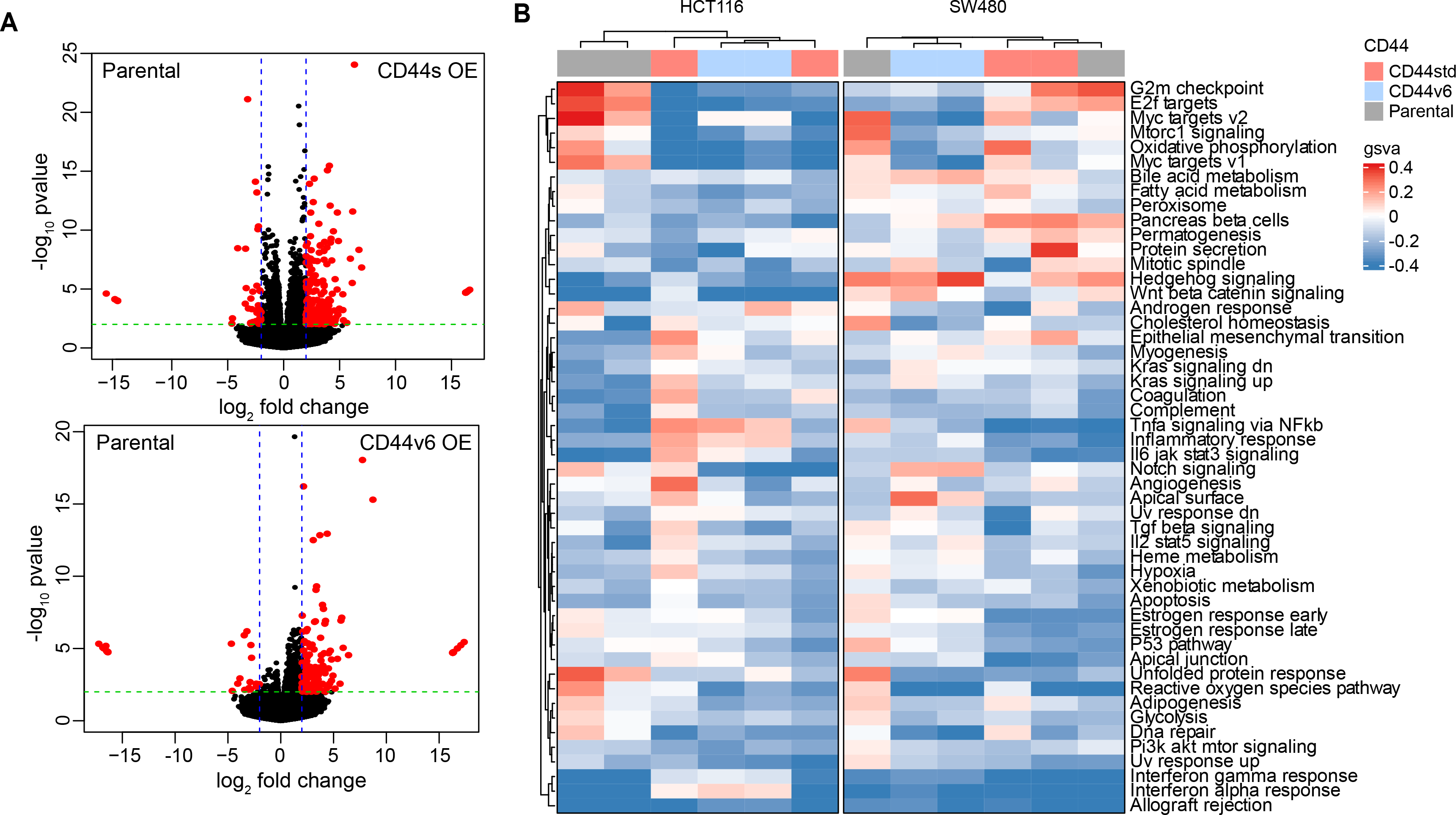
Gene Enrichment and Pathway Analysis of CD44s- and CD44v6-overexpressing colon cancer cells. **A.** Vulcano plots showing differentially expressed genes (absLFC > 2, pval < 0.01, red) by comparing parental cell lines to the CD44s- and CD44v6-OE samples in both cell lines. **B.** Gene Set Enrichment Analysis of parental, CD44s- and CD44v6-OE cells compared with each other in HCT116 and SW480 cells shown in heatmap, respectively. Only significantly altered pathways (NES > 1, and pval < 0.05) are shown.

**Figure S7.**
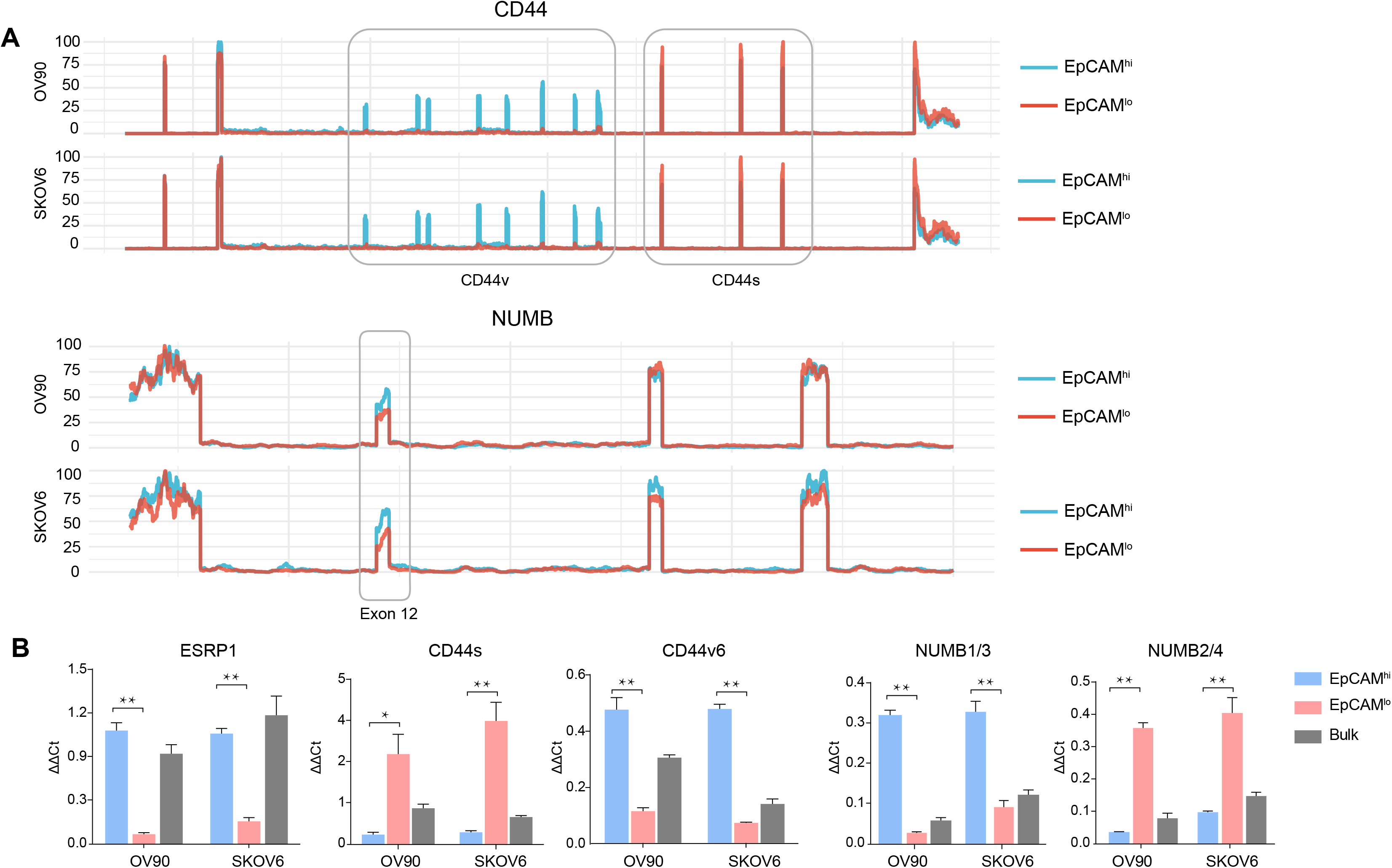
*CD44* and *NUMB* isoforms expression in EpCAM^hi/lo^ ovarian and cervical cancer cells. **A.** *CD44* and *NUMB* exon chromosome sites information from AS analysis in the ovarian and cervical cancer cell lines OV90 and SKOV6. Exon peak plot depicts the expression of different exons in the three groups; peak height is indicative of the expression level of specific exons. CD44v: CD44 exons v2 to v10. CD44v and CD44s, and NUMB exon 12 are highlighted by gray rectangles. **B.** RT-qPCR expression analysis of *ESRP1*, *CD44s*, *CD44v6*, *NUMB1*/*3*, and *NUMB2*/*4* isoforms in EpCAM^hi^, EpCAM^lo^, and bulk subpopulations in OV90 and SKOV6 ovarian cancer cell lines. *GAPDH* expression was used as control (Means±SEM, n=3). ** = p<0.01.

## Supplementary Table Legends

**Table S1.** List of alternative splicing targets in ESRP1 knocking down H358 line, HCT116 and SW480 EpCAM^lo^ and EpCAM^hi^ subpopulation, filtered by ΔPSI > 0.1

**Table S2.** List of alternative splicing targets in *ESRP1*-KD in the H358 cell line, *ESRP2*-KD in LNCaP, *RBM*47-KD in H358 line, QKI-KD in CAL27, and HCT116 and SW480 EpCAM^lo^ and EpCAM^hi^ subpopulation, filtered by ΔPSI > 0.1.

**Table S3.** Lists of primer sequences used for RT-PCR analysis.

**Table S4.** Differential expressed gene lists from the RNAseq analysis HCT116 CD44s- and CD44v6-OE cells.

**Table S5.** Differential expressed gene lists from the RNAseq analysis SW480 CD44s- and CD44v6-OE cells.

**Table S6.** List of Gene Set Enrichment Analysis (GSEA) in CD44s OE vs. CD44v6 OE vs. parental HCT116 and SW480 cells.

**Table S7.** List of Gene Set Variation Analysis (GSVA) in CD44s OE vs. CD44v6 OE vs. parental HCT116 and SW480 cells

## Notes

https://www.ncbi.nlm.nih.gov/geo/query/acc.cgi?acc=GSE192877

### Competing Interest Statement

The authors have declared no competing interest.

### Summary of Updates

Minor correction in the labeling of Supplementary Figures, in the text of the Materials and Methods and Results. Figure 2A-B, 2D, Figure 5C-D, Figure 4A-C, Figure6C, and Figure S3A-D have been modified.

